# PanClassif: Improving pan cancer classification of single cell RNA-seq gene expression data using machine learning

**DOI:** 10.1101/2021.04.10.439266

**Authors:** Kazi Ferdous Mahin, Md. Robiuddin, Mujahidul Islam, Shayed Ashraf, Farjana Yeasmin, Swakkhar Shatabda

## Abstract

Cancer is one of the major causes of human death per year. In recent years, cancer identification and classification using machine learning have gained momentum due to the availability of high throughput sequencing data. Using RNA-seq, cancer research is blooming day by day and new insights of cancer and related treatments are coming into light. In this paper, we propose PanClassif, a method that requires a very few and effective genes to detect cancer from RNA-seq data and is able to provide performance gain in several wide range machine learning classifiers. We have taken 22 types of cancer samples from The Cancer Genome Atlas (TCGA) having 8287 cancer samples and 680 normal samples. Firstly, PanClassif uses *k*-Nearest Neighbor (*k*-NN) smoothing to smooth the samples to handle noise in the data. Then effective genes are selected by Anova based test. For balancing the train data, PanClassif applies an oversampling method, SMOTE. We have performed comprehensive experiments on the datasets using several classification algorithms. Experimental results shows that PanClassif out-perform existing state-of-the-art methods available and shows consistent performance for two single cell RNA-seq datasets taken from Gene Expression Omnibus (GEO). PanClassif improves performances of a wide variety of classifiers for both binary cancer prediction and multi-class cancer classification. PanClassif is available as a python package (https://pypi.org/project/panclassif/). All the source code and materials of PanClassif are available at https://github.com/Zwei-inc/panclassif.

## 1 Introduction

Cancer remains one of the most spiteful diseases that tribulates human life. It comprises a convoluted biological system that requires meticulous and all-inclusive analysis. Over the last couple of decades, RNA sequencing has become a remarkable way for transcriptome profiling. The revolution sparkled when high throughput sequencing evolved from bulk RNA sequencing to single molecular, single cell [1]. Spatial transcriptome analysis methods have enabled increasingly meticulous cell resolution incorporated with spatial information [2].

Heterogeneity of cells which comes from different kind of gene regulation, gene mutations and stochastic dissimilarity are reflected at the transcriptomic, proteomic and genomic levels. This kind of heterogeneity is a significant factor in cancer treatment failure. Cancer is itself a complex disease but during formation of malignant cells, the lineages divide and form intra-tumour heterogeneity [3]. It is clear that conventional bulk detection method which measures the average profile of the tumour population may have limitations to characterize intricate diseases like cancer. In the case of single cell RNA-seq experiments, challenges in measuring technologies like amplification bias, library size differences, cell cycle and low capture rate of RNA lead to substantial noise in which is known as dropouts [4]. It necessitates different smoothing techniques to be applied to the data.

In cancer classification, it is crucial to identify features explicitly because of the high dimensionality of the data. In recent years, machine learning based methods have gained popularity in cancer cell classification due to the effectiveness in identifying crucial features and availability of data from high throughput machines [5].

Li et al. [6] have used RNA-seq expression of 9096 tumor samples which represent 31 types of tumor data collected from TCGA. They have used genetic algorithms for gene selection, *log*_2_-transform for normalizing data and *k*-NN for classification. After selecting 20 genes, they have performed a classification experiment with those data sets. They have also found several genes that are responsible for showing sexual-dimorphism in liver cancer. Hsu et al. [7] takes 33 types of cancer from TCGA and does a comparison of different machine learning models based on prediction accuracy, recall, precision, *F* -score, and training time. The models they have chosen are Decision Tree, *k*-NN, *linear* and *poly* kernel Support Vector Machine (SVM), and Neural Networks (NN).

Lyu et al. [8], have used 33 types of cancer data collected from TCGA. After performing log2 transform on the whole dataset, they have reduced 1000 unannoted genes which were not found in NCBI gene annotation file. Then, they have removed the genes having variance across samples less than 1.19 and left with 10381 genes. After that, they have made each sample into a 102×102 2-D image and used Convolutional Neural Network(CNN) with 3 layers to perform classification.

Kim et al. [9] have classified 21 types of cancer using single-cell RNA-Seq gene count data which is collected from TCGA and then filter out genes from each cancer based on the highest *F* -score. They have chosen *n*-numbers of genes for training features and figures out the best *n* genes which gives the best performance. After that they have used *z*-score for normalizing the data and perform binary and pan-cancer classification using machine learning algorithms. They have used breast cancer and skin melanoma cancer dataset from Gene Expression Omnibus (GEO) repository to validate the overall performance of their model.

In [10], the authors have used RNA-seq gene count data and filter out genes that have zero counts in more than 80% of samples. Then they have augmented the data to eradicate the effect of noise in training. Their augmented approach that has made data scale-invariant and finally classify data with the neural network. Guo et al. [11] have discussed a bout molecular heterogeneities and complex micro-environments which bring great challenges for diagnosis of cancer as well as treatment procedure with the help of RNA Seq. Focusing on the process and analysis of single cell RNA-seq data for cancer research, they have developed an R package called *scCancer* which has integrated the basic steps of processing single cell RNA-seq data. It has used common batch-effect correction methods for performing multi-sample integration analysis. They propose a new algorithm named ‘NormalMNN’. It can automatically generate user-friendly graphic reports to overview all the results. The experimental dataset that they used for testing has 433405 cells from 56 samples.

In [12], the authors have proposed SCOPE (Supervised Cancer Origin Prediction Using Expression) where they have used TCGA data for training their machine learning model and used untreated primary mesothelioma male, treatment-resistant cancer female patients data for testing. They have taken the common genes among the training and testing data. They have not performed any feature selection technique, thus used all the features for training their models. They have used SVM, Random Forest, Extra trees and Fully connected neural network as the classifiers of their model. They have used total 66 classes (40 cancer, 26 normal) for training the model.

Transcriptional signature can be used to classify cell types according to its transcriptional profile. Single cell RNA-seq technique appeared to be one of the most reliable technique in finding unique genes in a cell. Alquicira-Hernandez et al. [13] have proposed a method named ‘scPred’ which can confidently predict accurate single cell states using their transcriptional profile. ScPred splits its model into training step and prediction step. In training step, using singular value decomposition a gene expression matrix representation has been made. After going through the feature selection step, a SVM model has been fitted. In prediction step, informative principal components are used to predict the class probabilities.

Mercatelli et al. [14] have predicted pancancer classification and relation between somatic mutation and gene expression of single cell data. They have used 24 TCGA dataset to predict the somatic mutation and copy the number of variations to different cancer cells. Then finally, using ARACNe/VIPER or WGCNA methods to clustering the group of expressed genes. Ultimately. their proposed principle can have a good impact in noisy datasets based on scRNA-Seq.

Ferreira et al. [15] have compared between three Auto Encoders(AE) where, the AEs are used as deep neural network weight initialization technique. The three AEs are basic AE, denoising AE and sparse AE. They have followed two approaches for training the classification networks and another two approaches for embedding the AEs in the classification network. For their experiment they have used 5 different cancer type (thyroid, skin, stomach, breast, and lung) which were obtained from TCGA. And for validating their experiment pipeline, they have used Malaria and Wisconsin Breast cancer data.

Khalifa et al. [16] have proposed an optimized deep learning approach where, BPSO-DT(Binary Particle Swarm Optimization with Decision Tree) used for feature selection and CNN (Convolution Neural Network) for classifying the different type of cancer. They have used 5 cancer dataset (KIRC, BRCA, LUSC, LUAD, UCEC) for their work. Among 971 features, BPSO-DT have chosen 615 best RNA-seq feature. Then, they have scaled the data in [0-255] and then converted each cancer sample into 25×25 2-D image. After that, they have performed augmentation on the whole data. Finally, they have used CNN for the classification.

Hou et al. [17] have evaluated the impact of different single-cell RNA sequencing imputation approaches. Many single-cell methods for reducing systematic technical noises have been presented in recent years, including imputation methods, which try to address the increased sparsity observed in single-cell data. Despite the fact that several imputation methods have been developed, there is no agreement on how they have compared to one another. They have evaluated 18 single-cell RNA seq imputation methods to see how accurate and usable they were. In bulk RNA-seq, the majority of scRNA-seq imputation approaches outperformed no imputation in recovering gene expression. However, the majority of the approaches, particularly clustering and trajectory analysis, did not increase performance in downstream studies when compared to no imputation, and should thus be utilized with caution. Overall, MAGIC [18], kNN-smoothing [19], and SAVER [20] were found to consistently outperform the other methods.

Single cell genomics pave the way of identifying unique traits and genes in individual cells. To gain more biological insights from single cell data, it is better to have the data annotated according to cell identity. But manual annotation may arise errors and consume more valuable time. Jacob et al. [21] have come with the idea of transferring cell identity information among experiments using adversarial neural network model. This neural network are taught the representation of the cell identity. Simultaneously, an adversarial model estimates a domain of cell from taught embedding. ‘scNym’ confidently shares annotation among all the experiments in spite of biological and technical variances.

In spite of having supremacy in cancer cell classification, single cell RNA-seq bears the curse of dimensionality. To eradicate this curse. Ge et al. [22] has proposed a domain adversarial neural network model. Generally dimensionality reduction arise another issue called batch effect which refers batch representation rather than cell type specific representation. Thus fails to locate significant genes. To prevent batch effect, their model continuously evaluates the accuracy of cell type assignment and enhances the inability to distinguish the batch.

In cell based studies, unsupervised methods are commonly used. Due to unsupervised clustering, most of the existing methods may not produce biologically interpretable clusters. The main reason behind the problem is high dimensionality of scRNA-seq data and dropouts. Unlike traditional methods, Tian et al. [23] has initiated a deep learning model which learns simultaneously and also updates loss function with soft pairwise constraints. This soft pairwise function are from converted prior knowledge.

Single-cell RNA Seq is the next-generation technology that is able to detect cancer more accurately. However, the identification of different cancer cells from a multi-cancer data sample is still challenging. Single cell RNA-Seq datasets comes with a huge number of features or gene expressions and as well as with noise. As these datasets have high dimensionality, machine learning models often take too much time and resource to train and perform prediction. In addition, there is an imbalance in the datasets. Often the ratio of normal and cancer cell is not balanced as shown in Table 1 and Table 2. Training with imbalanced data leads to biased models with lower sensitivity.

**Table 1:** Breast cancer and skin melanoma cancer datasets from GEO.

| **Cancer Type** | **Accession code** | **Cancer** | **Normal** |
| --- | --- | --- | --- |
| Breast Cancer | GSE75688 | 317 | 198 |
| Skin Melanoma | GSE72056 | 1257 | 3256 |
| **Total** |  | 1574 | 3454 |

**Table 2:** TCGA cancer codes and their respective cancer and normal samples used in our work.

| **Code** | **Cancer** | **Normal** | **Code** | **Cancer** | **Normal** |
| --- | --- | --- | --- | --- | --- |
| BLCA | 427 | 19 | LUAD | 576 | 58 |
| BRCA | 1212 | 112 | LUSC | 552 | 51 |
| CESC | 309 | 3 | PAAD | 183 | 4 |
| CHOL | 45 | 9 | PCPG | 187 | 3 |
| COAD | 328 | 26 | PRAD | 550 | 52 |
| ESCA | 198 | 11 | READ | 105 | 6 |
| HNSC | 566 | 43 | SARC | 265 | 3 |
| KICH | 91 | 25 | STAD | 450 | 32 |
| KIRC | 606 | 72 | THCA | 568 | 59 |
| KIRP | 323 | 32 | THYM | 122 | 3 |
| LIHC | 423 | 50 | UCEC | 201 | 7 |
| **Total** |  |  |  | 8287 | 680 |

In this paper, we propose PanClassif to address all the above mentioned issues. PanClassif requires a very small amount of genes to correctly classify different types of cancer samples with a very few miss-classifications. We have taken 22 types of cancer data from TCGA. After that we have used smoothing on the datasets using *k*-NN smoothing and then performed ANOVA based gene selection. After picking the best genes we filter out rest of the genes from the dataset. We then perform oversampling to handle the data imbalance problem on the training data. Then we evaluate the performance of the method using several machine learning models. Most of the models perform very well. We have chosen 6 classifiers: SVM (*linear* kernel), SVM (*rbf* kernel), Random Forest, Neural Network, *k*-Nearest Neighbour and Adaboost algorithm. Most of the classification algorithms have shown increase in performance following PanClassif methodology. We further performed gene set enrichment analysis over the best selected genes from TCGA cancer datasets and most of the selected genes have shown correlation with cancer/tumor type diseases. Experiments show the effectiveness of the method on Gene Expression Omnibus (GEO) single-cell RNA-Seq datasets. The results obtained by PanClassif are significantly improved over existing state-of-the-art methods.

## 2 Materials and Methods

In Fig. 1, we have shown the complete workflow of PanClassif. At first, we have downloaded our desired data from TCGA and GEO and then PanClassif applies *k*-NN smoothing on our downloaded data. Then feature selection is performed to select an effective set of features followed by the standardization of the data. After standardization, PanClassif splits the data in train and test. Oversampling is performed on the train data to address the imbalance issue. Finally, PanClassif trains the data for binary and multiclass classification models using training data and validate the model’s performance using test data. In this paper, two classification problems are addressed: binary and multi-class. In binary classification, we checked if a sample has cancer or not and in multi-class classification, we checked which type of cancer a sample has. The detailed discussion about each component of PanClassif are described in the rest of the section.

**Figure 1:**
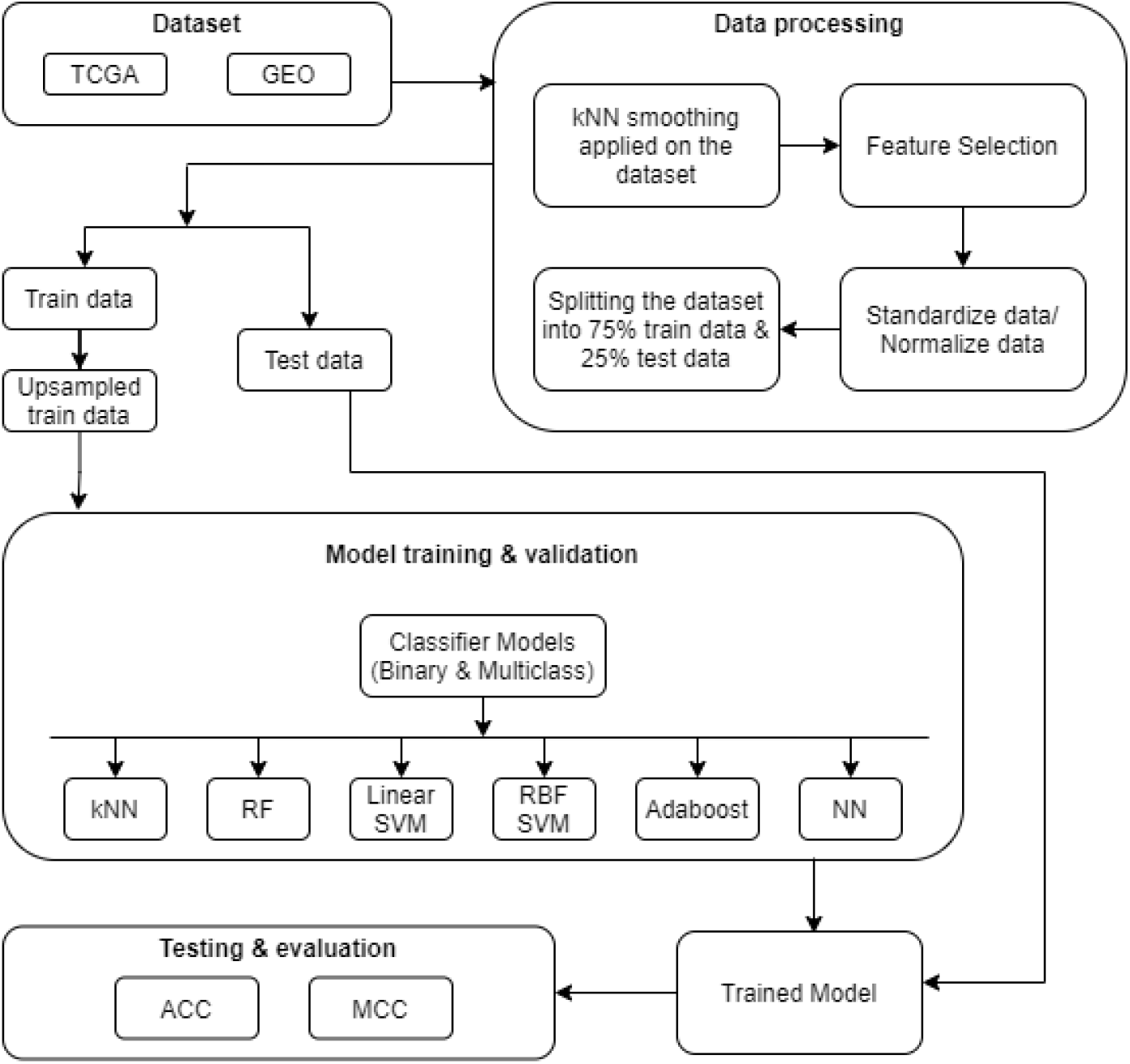
Complete workflow of PanClassif.

### 2.1 Data Collection

In this paper, we have used RNA-seq expression count data of cancerous and their respective normal for this research. We have collected data from the repositories of The Cancer Genome Atlas (TCGA) and Gene Expression Omnibus (GEO). From TCGA, we have collected samples of 22 types of cancer and from GEO, we have collected breast cancer (GSE75688) [24] and skin melanoma cancer (GSE72056) [25] samples. The expression counts were calculated using RSEM [26] on the data downloaded using TCGA2STAT [27]. TCGA2STAT is also used to download TCGA data. Total 8287 cancer samples and 680 normal samples are collected from TCGA having 20,501 genes. Breast cancer (BRCA) and skin melanoma cancer (SKCM) data that are collected from GEO were also calculated using RSEM and have 317 cancer samples, 198 normal samples and 1257 cancer samples, 3256 normal samples respectively. Datasets collected from both TCGA and GEO are not log-transformed. The summary of the datasets collected are given in Table 1 and Table 2. Both of the datasets were then prepared for binary cancer prediction and multiclass cancer type identification problems. Single cell RNA-seq data from GEO datasets were used to validate the effectiveness of the methodology.

### 2.2 Data Smoothing

We have already mentioned that single-cell RNA-seq data tends to be extremely noisy. In order to remove the effect of noise and variability of the data, we have used *k*-nearest neighbour smoothing [19], SAVER [20].

The *k*-NN smoothing uses step-wise approach to identify the nearest neighbours and Freeman-Tukey transformation to ensure that in the distance calculation, all genes expression levels contribution are approximately equal. In addition to this, Principal Component Analysis (PCA) is also used to remove noise and give attention to the most important expression differences among cells. We have performed this smoothing on our data samples using *k* = 10. Dataset having less than 10 samples is skipped in this process.

According to [20], based on single-cell RNA sequence data, they suggested an imputation method for UMI (Unique Molecule Index). Only a small fraction of the transcripts present in each cell are sequenced in single-cell RNA sequencing studies. This results in erroneous quantification of genes with low or moderate expression, which obstructs down-stream analysis. SAVER (Single-cell Analysis Via Expression Recovery) is their proposed method for recovering the true gene expression profile in noisy and sparse single-cell RNA-seq data. It borrows information across genes and cells and uses regularized regression prediction and the empirical Bayes method to recover the true gene expression profile in noisy and sparse single-cell RNA-seq data.

Among this two methods *k*-NN smoothing outperforms SAVER so we choose *k*-NN smoothing for our proposed method.(See supplementary table S16.)

### 2.3 Feature Selection

PanClassif uses *F* 1-Anova test to select effective set of genes from the huge number of genes for which the expression data is available. For identifying the best genes in our paper, we utilized one-way ANOVA. ANOVA is used to calculate the dispersion of gene counts in genes. One-way ANOVA enables a gene to compare the means of two or more groups of genes and choose the best genes. If a gene’s variation is modest, its influence on the dataset is also minimal. We utilize *SelectKBest* on those anova values since this technique only selects the top K genes based on anova value, which implies it selects the best impacted genes from each dataset. The one-way ANOVA hypothesis examines the null hypothesis that two or more groups have the same population mean. The test is performed on samples drawn from two or more groups, which maybe of varying sizes. The feature selection algorithm is depicted in Algorithm 1. For each cancer type, first the dataset for the particular cancer is extracted from database as gene expression count. Then Anova test is performed to get the scores. Top *n* genes are selected and only the unique genes are added to the selected gene list. Here, in our experiment the value of *n* is 5, 10, 30, 40, 50 and get 105, 204, 571, 743, 899 unique genes respectively for TCGA data. But for GEO, since there are only three class so to match our result with TCGA’s data we have used *n* = 105, 204, 571, 743, 899. We have performed detailed experiments to show the effectiveness of this feature selection algorithm and the effect of the parameter *n*. The algorithm is initially experimented on TCGA dataset and then validated using the same parameters in the GEO dataset.

We have provided some step-by-step equations for how F1-ANOVA generates F-Score for each feature. Here, *SSB* = sum of square between the groups, *SSE*_*j*_ = sum of squares due to error for *j*^*th*^ feature, *j* = feature index, *n*_*j*_ = list of total samples per feature, 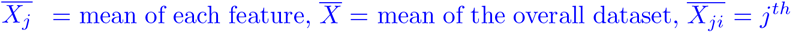 feature’s all samples, *MSW*_*j*_ = mean sum of square within the group, *MSB* = mean sum of square between the groups, *F*_*j*_ = f-score for *j*^*th*^ feature. When we use the *SelectKBest* it executes these equations sequentially for each feature and return the *k* best features based on this f-score.

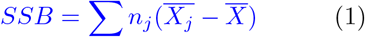

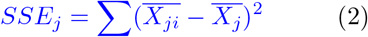

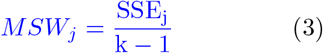

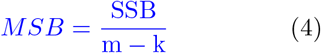

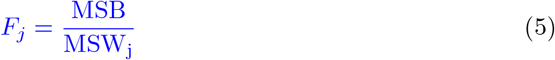

### 2.4 Standardization and Sampling

Feature scaling or standardization is a very important step in data pre-processing. It helps to reduce effect of any gene that shows high expression count and can effect the classification model performances. PanClassif uses *z*-score standardization on the datasets. Standard score of a sample *x*_*ij*_ drawn from a population *j* with mean, *µ* can be calculated

#### Algorithm 1: Feature Selection

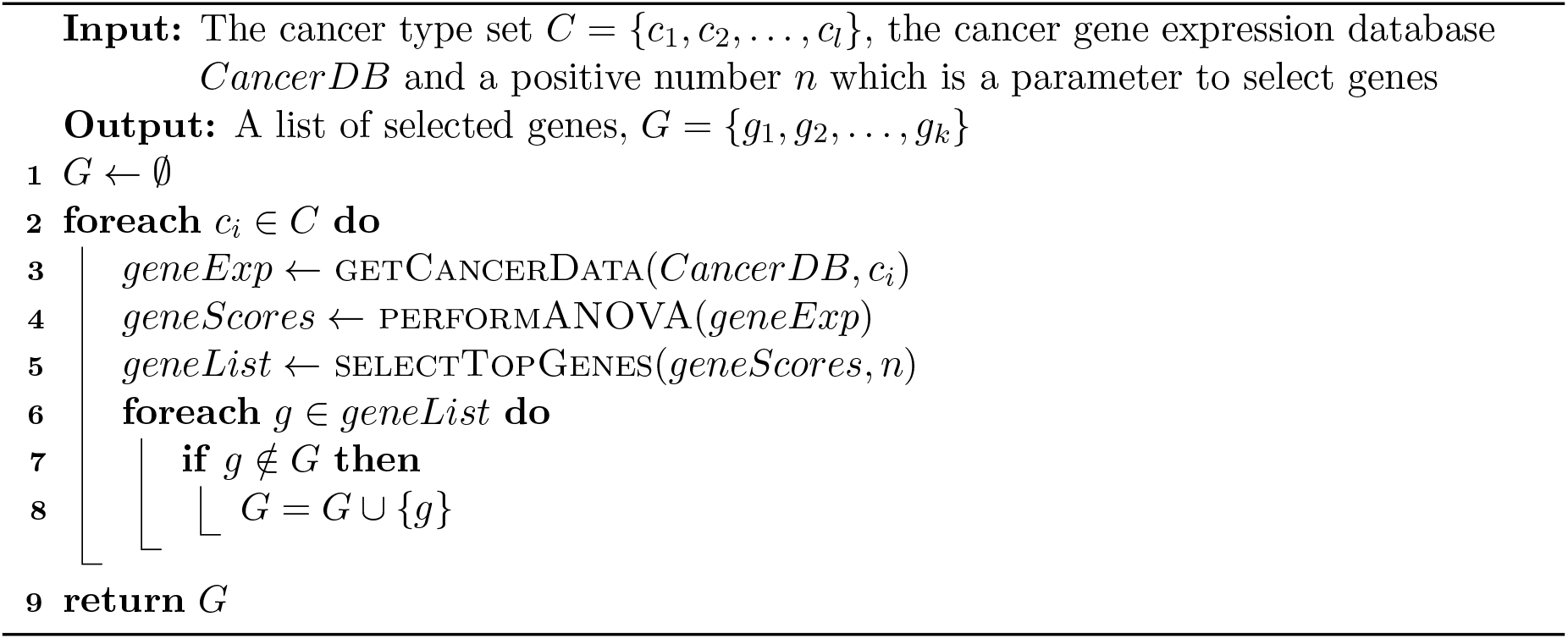

using:

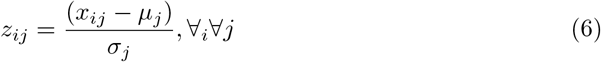

Here, *σ*_*j*_ is the standard deviation of the population. Here, *µ*_*j*_, *σ*_*j*_ are estimated using the sample mean and standard deviation.

After standardization, data was randomly split into train and test datasets with a 3:1 ratio. We have used this ratio as done by [9]. In both of the datasets, we could notice that there is an imbalance in the number of positive and negative classification. particularly for the TCGA datasets, the number of normal or non-cancer data is very low. In case of binary classification the ratio is 12:1. The GEO dataset is also imbalanced in the other way around. Such imbalance creates a bias in the learned model. To address this problem, PanClassif uses Synthetic Minority Oversampling Technique (SMOTE) [28] to oversample the train dataset by generating synthetic data in the minority class.

### 2.5 Machine Learning models for classification

The selection of proper classification model is a very crucial step in machine learning. However, to show the effectiveness of PanClassif, we have selected a number of existing classifiers to apply them for binary and multi-class classification tasks. They are *k*-Nearest Neighbour (*k*-NN), Support Vector Machines with linear kernel (L-SVM), Support Vector Machines with radial basis function kernel (RBF-SVM) [29], Random Forest (RF) [30], Adaboost [31] and Artificial Neural Networks (NN) [32]. In *k*-NN, we have used *k* = {1, 2, 3, 5, 10, 13, 17}. For RF, the *maxDepth* parameter of decision trees were kept 100. For both of L-SVM and RBF-SVM, *C* was kept 1 and *γ* was set at the inverse value of multiplication of no. of the features in the dataset and their variance. Decision tree is used as base classifier for Adaboost. For neural network, we have used 5 hidden layers apart from the input layer and output layer. The number of neurons in the hidden layers were 512, 256, 128, 64, and 32 respectively. Same hyper-parameters are used in both binary and multi-class classification tasks except in the neural network. In neural network, sigmoid is used as last layer activation for binary and softmax is used as last layer activation in multi-class classification.

### 2.6 Performance Evaluation

For evaluating the performance of the classification models and overall PanClassif methodology we have used two metrics: Accuracy (*ACC*) and Mathews Correlation Coefficient (*MCC*). In the case of binary classification task, true positives (*TP*) is the number of positive class instances classified correctly and true negatives (*TN*) are the number of negative class instances classified correctly by the model. On the other had false positives (*FP*) and false negatives (*FN*) are number of negative and positive class instances respectively that are wrongly classified in the opposite group. Based on these, accuracy and MCC are defined as in the following.

#### Accuracy

Accuracy determines how accurately the model classifies the instances. It has a range of 0 to 1 where a value 1 indicates the best possible classifier. However, please note that for imbalanced data, accuracy might not always reflect the proper model performances. Accuracy is define in the following equation:

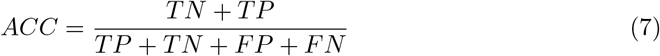

#### Mathews Correlation Coefficient

Mathews Correlation Coefficient or MCC provides a good performance measure even when the data is imbalanced. It has a range of -1 to +1. A perfect model will show a value close to +1 and a less accurate model will achieve a value less than that and even negative.

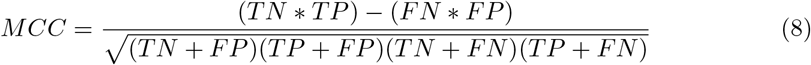

Calculating accuracy and MCC score for multi-class is different than binary. In the case of multi-class task, we have used a *one vs all* for each class and then calculated accuracy and MCC score for each class. After that, the weighted average value of each metric is reported.

## 3 Results and Discussion

All the experiments done in this paper and the reported results are prepared using a machine having 32GB RAM, Xenon processor, and Linux operating system. All the programs are written in Python 3.8 and Scikit-learn library. Each experiments were run 5 times and only the average results are reported. Please note that Supplementary data contains more detailed results on the datasets and methods used in this paper.

### 3.1 Performance on TCGA data

For both binary and pan-cancer classification, we have used 75% of the data for training and the rest of the data for test purpose. In case of binary classification task with TCGA data, the training data has a very high imbalance ratio of 12:1 with 6213 cancer samples and 510 normal samples. The oversampling techniques reduced this ratio to 3:1 resulting into 2057 normal samples. We have annotated two classes, ‘cancer’ samples as class 1 and ‘normal’ samples as class 0. Then we have prepared 5 different datasets with top 105, 204, 571, 743, 899 genes (keeping the parameter in feature selection *n* = 5, 10, 30, 40, 50 respectively) and train models with those 5 datasets and test with the remaining 25% test data. A spider plot of accuracy score achieved by different classifiers on different number of selected genes are shown in Fig. 2. From the results shown in Fig. 2, we observe that *k*-NN and Random Forest (RF) are the two best performing classifiers. Adaboost (ADB) and the variants of SVM did not perform well in any of the datasets compared to them. However, performance of Neural Network was very close to the best results where number of selected genes were 204, 571 and 899. (See supplementary table S1 for details.) Please note that, results in terms of MCC also shows superiority of Random Forest and *k*-NN over other classifiers. (See supplementary figure S1.)

**Figure 2:**
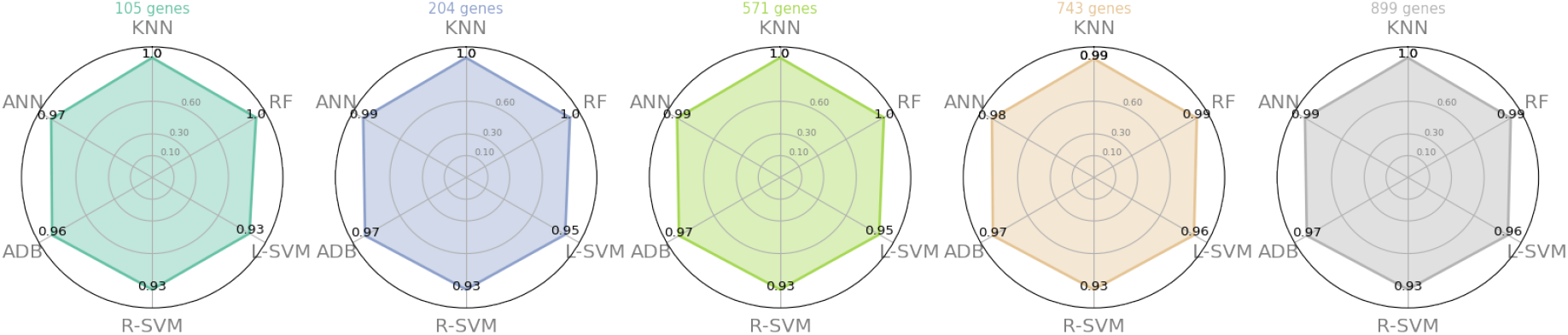
Spider plot for ACC score for the binary classification task in TCGA data using 5 different datasets created selecting different number of genes.

For pan cancer classification, we create 5 datasets with top 105, 204, 571, 743, 899 genes in the similar way and annotate them with 23 classes where 22 types were of cancer class and 1 normal class. After that, we have trained all 6 Machine Learning models with the same configurations described in Section 2.5. The train and test set ratio was kept similar as in the binary prediction task (3:1). A spider plot of accuracy scores achieved by the different classifiers using different number of genes is shown in Fig. 3. The plot shows a very poor performance by the AdaBoost algorithm. The SVM classifier with radial basis function as kernel showed some satisfactory results. However, the rest of the classifiers performed very well in terms of accuracy (for results in terms of MCC, please see Supplementary Figure S2 and Table S3). Among all the classifiers, Random forest performed the best and the confusion matrix for Random forest in the multi-class classification is shown in Fig. 6. From this matrix, it is observable that there were no inter-class mistakes done by the classifier. However, a few false positives and false negatives were found for cancer types BRCA, BLCA, KICH, LIHC, KIRP, THCA.

**Figure 3:**
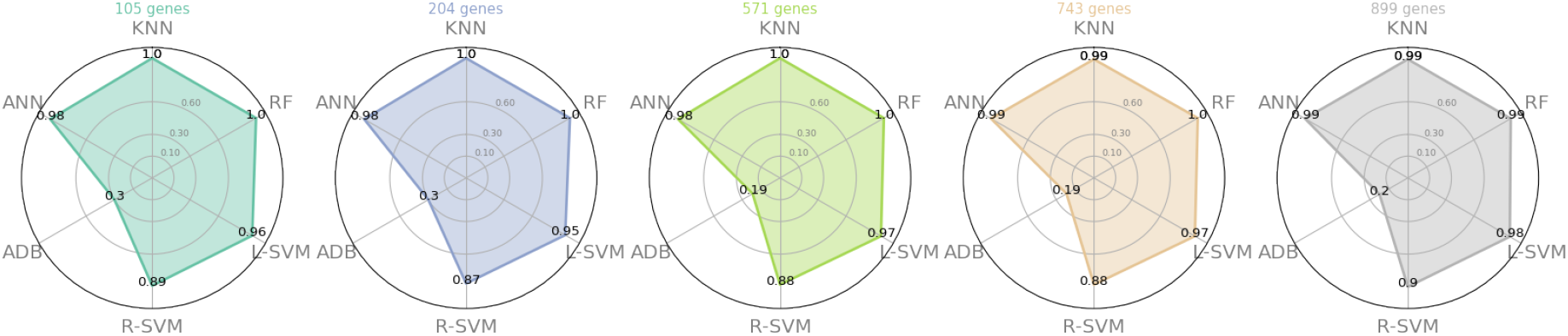
Spider plot for ACC score for the multi-class classification task in TCGA data using 5 different datasets created selecting different number of genes.

From Figure 2 and Figure 3, we can observe that the performances for best genes set from 105-899 are almost similar in terms of ACC. Since we are dealing with imbalanced datasets of single-cell RNA-seq, accuracy does not indicate much significance. Therefore, we are using MCC to figure out the best genes set for our classifiers. From supplementary Figure S1 and S2, we can easily determine the changes of result in different classifiers using different genes sets.

### 3.2 Performance on GEO data

To validate the performance of PanClassif, we have used single cell RNA-seq GEO data. The setting and parameter of PanClassif pipeline were kept same except the value *n*. Since GEO’s data contains only three class. So, we have chosen *n* = 105, 204, 571, 743, 899 to match with number of unique genes in TCGA. Similar result pattern is observed here too. Figure 4 and 5 shows that MCC score of Random Forest classifier is similar to TCGA in both binary and multi-class classification tasks. Instead of *k*NN, Adaboost is performing better here. As mentioned before, accuracy metric does help to distinguish the best classifier nor best genes set in case of imbalanced data. Here from Figure 4 and Figure 5, we can see the changes in MCC score of different classifiers in different genes set are more evident. In figure 4, we observe MCC scire of *k*NN and R-SVM decreases as we increase the value of *n*. On the other hand, MCC of L-SVM increases as value of *n* increases. RF, ANN and ADB shows consistency in MCC score. In Figure 5, L-SVM performance was getting worse as the *n* value increases. MCC score of *k*NN and R-SVM decreases as we increase *n*’s value. ANN, RF and ADB’s shows consistent performance but in 899 genes set, ADB’s performance decrease (See supplementary Table S2 and Table S4 for details).

**Figure 4:**
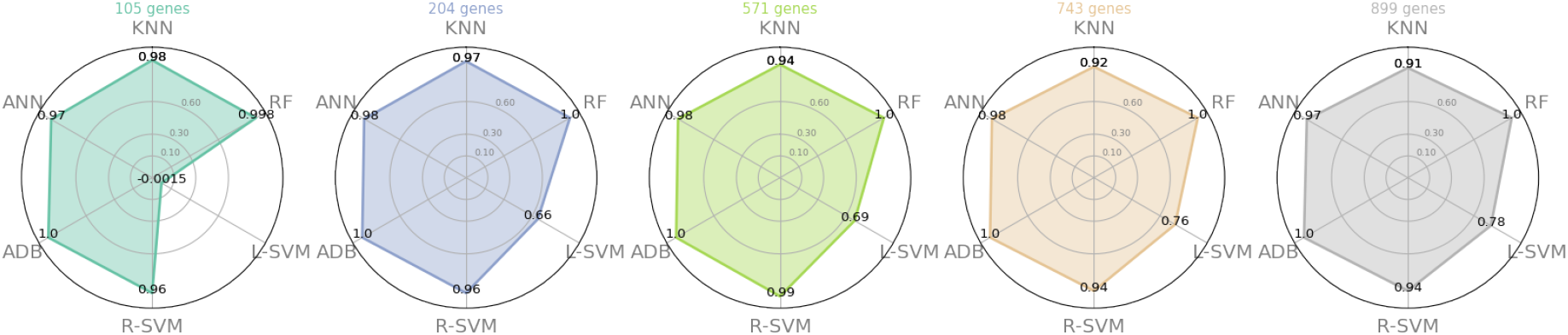
Spider plot for MCC score for the binary-class classification task in GEO data using 5 different datasets created selecting different number of genes.

**Figure 5:**
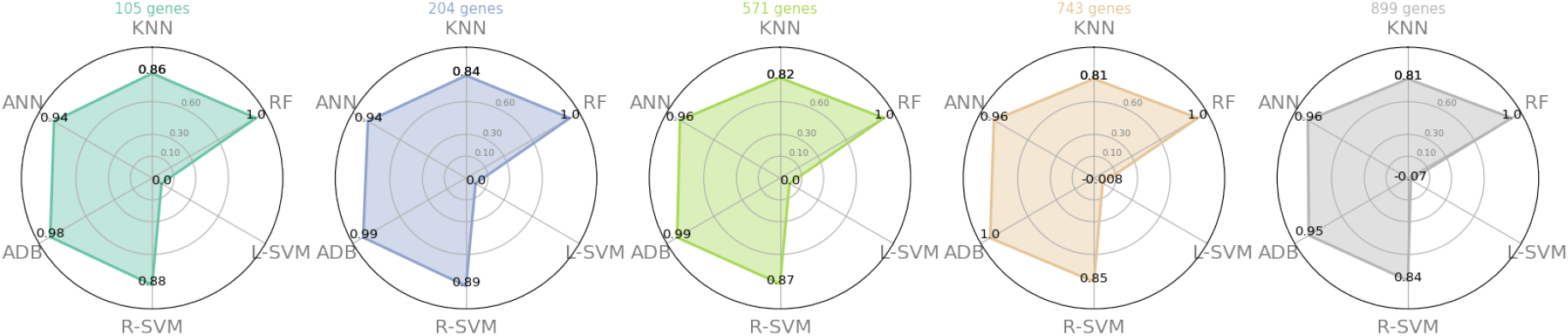
Spider plot for MCC score for the multi-class classification task in GEO data using 5 different datasets created selecting different number of genes.

**Figure 6:**
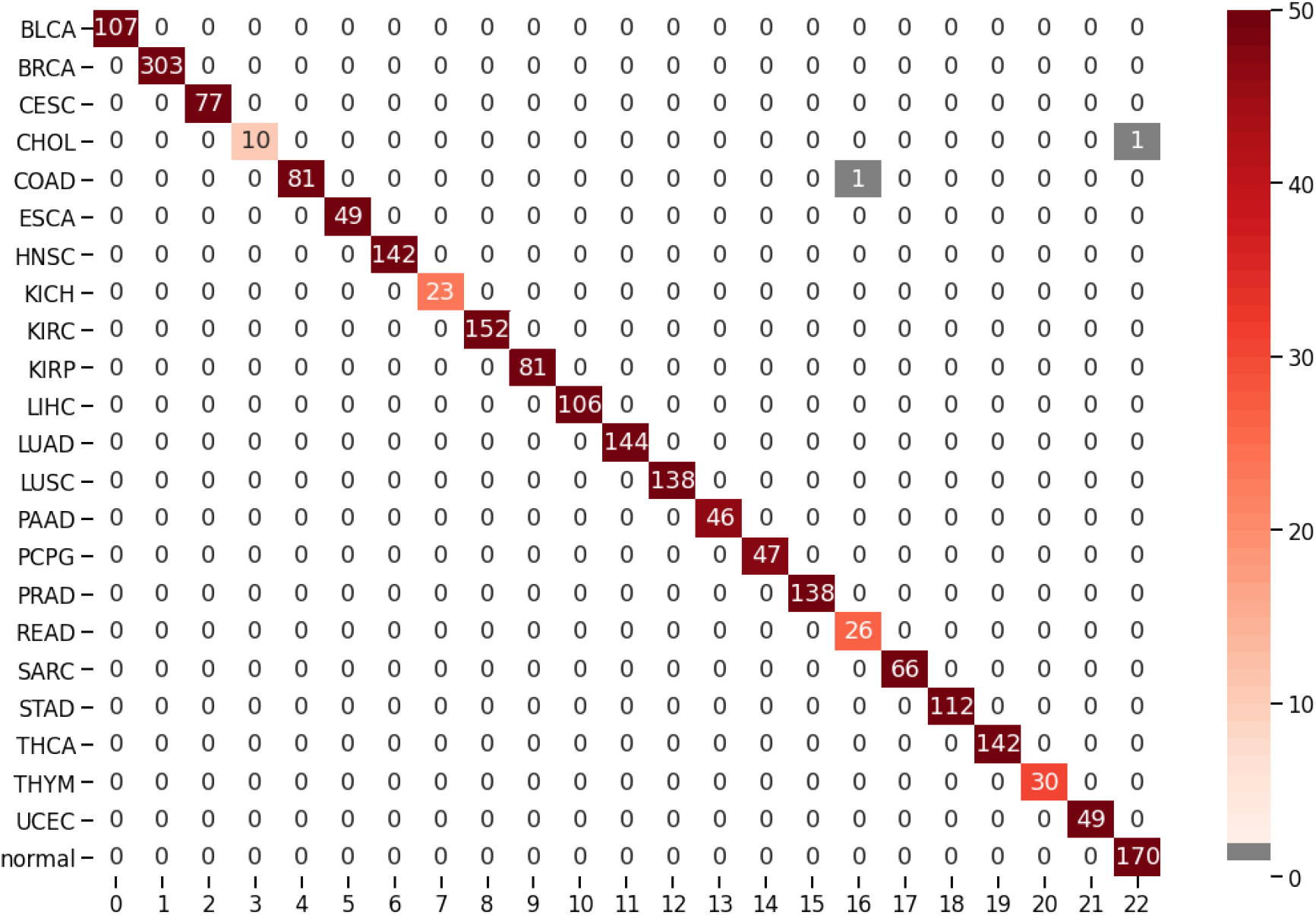
Confusion matrix of multi-class classification by Random Forest Classifier.

### 3.3 Comparison with other methods

We have also performed a comparative analysis for the performance of PanClassif with that of the work in [9], [8], [6], [15], [16] which is a state-of-the-art method for cancer classification and recognition task from single cell RNA-Seq data. Table 3 shows the MCC and accuracy achieved by these methods on TCGA data for both of the tasks using different classifiers. The results reported in the table were taken from the reported results by [9]. For binary classification task, [9] achieved highest accuracy of 0.99 with 300 genes using neural network. However, using PanClassif, accuracy of neural network is 0.99 using 204 genes. Our best classifier is random forest which gives 1.00 accuracy with 105 genes. In pan-cancer classification the performance of PanClassif remain same. However, for pan cancer classification, [9] achieves a best accuracy of 0.90 by neural network using 300 genes. PanClassif achieves 0.98 accuracy using neural network with 105 genes. [9] achieved best performances using in between 200 to 300 genes for binary and pan-cancer classification. In our case, the range of genes is 105-204 in most cases except SVM with *rbf* kernel shows top-notch performance. In case of MCC score, PanClassif also significantly outperform state-of-the-art method.

**Table 3:**
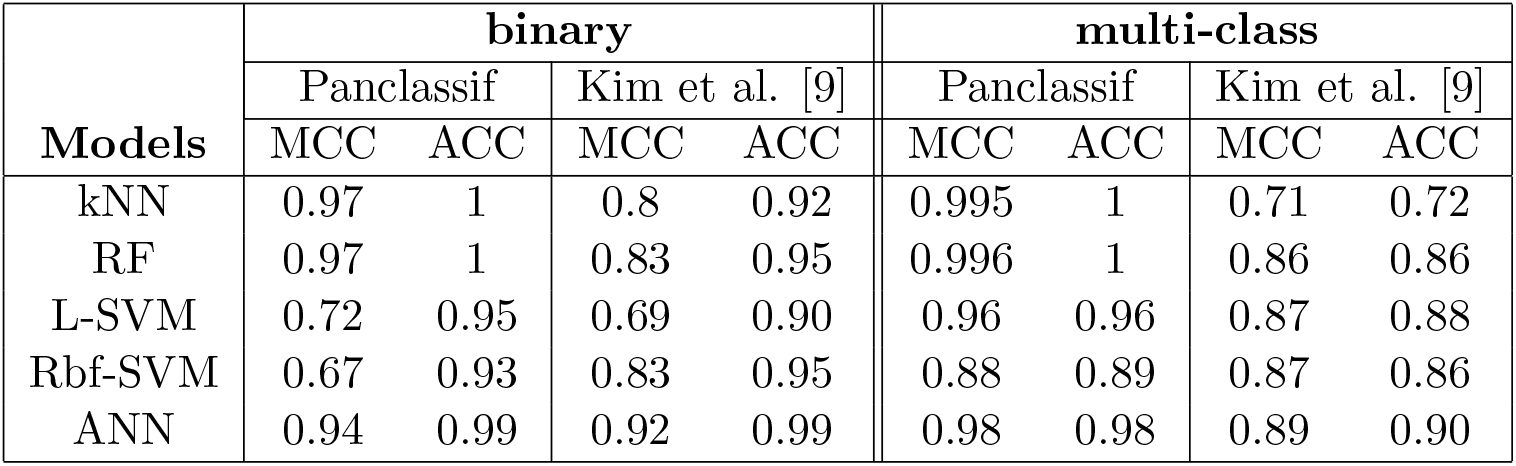
Performance comparison of PanClassif with Kim et al. [9] on TCGA dataset.

In case of the GEO datasets, [9] achieves best MCC of 0.85 for binary classification and 0.86 in multiclass classification as reported in their paper. Their best model is ANN and 0.85 and 0.86 are the MCC score of ANN classifier. For PanClassif, ANN got 0.97 MCC score in binary and 0.93 in multiclass classification. Except for SVM with *linear* kernel, all of our classifiers using PanClassif get significantly higher MCC score than their best reported model.

In Table 4, other works that addressed the cancer classification task using TCGA dataset are compared with PanClassif in terms of accuracy. From the Table 4, we observe that in the case kidney adjacent classes (KICH, KIRP, KIRC) some misclassifications are there for the methods proposed in [8] and [6]. Also, we can note while predicting CHOL and READ [8], [6] have performed significantly worse compared to PanClassif. Now from the table mentioned above, it is clearly distinguishable that PanClassif has outperformed the other methods. Please note we have reported the accuracy results as it has been presented by the authors in their respective papers.

**Table 4:**
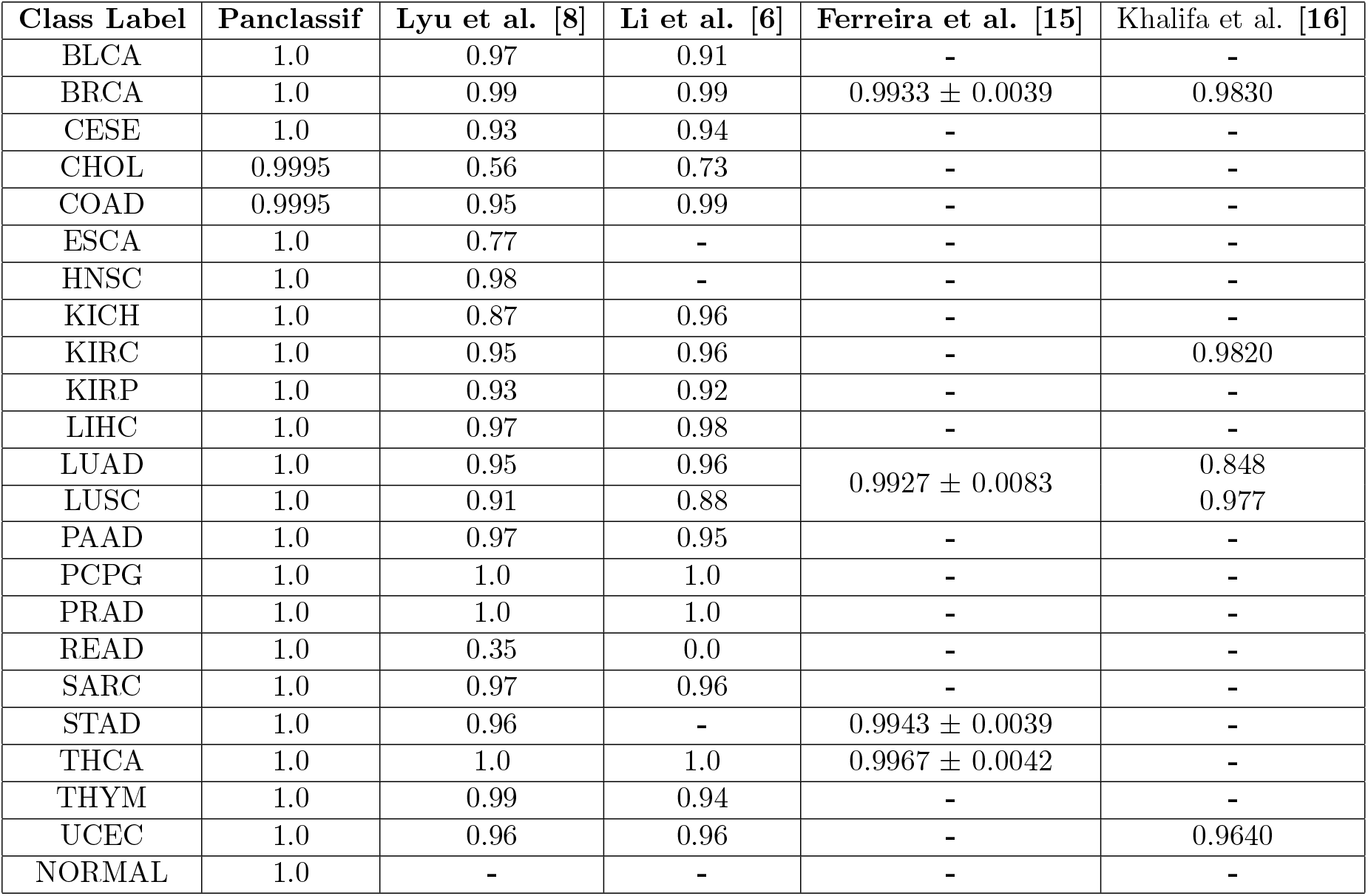
Performance comparison of PanClassif with [8], [6], [15] and [16] on TCGA dataset using accuracy.

| Class Label | Panclassif | Lyu et al. [8] | Li et al. [6] | Ferreira et al. [15] | Khalifa et al. [16] |
| --- | --- | --- | --- | --- | --- |
| BLCA | 1.0 | 0.97 | 0.91 | - | - |
| BRCA | 1.0 | 0.99 | 0.99 | $0.9933 \pm 0.0039$ | 0.9830 |
| CESE | 1.0 | 0.93 | 0.94 | - | - |
| CHOL | 0.9995 | 0.56 | 0.73 | - | - |
| COAD | 0.9995 | 0.95 | 0.99 | - | - |
| ESCA | 1.0 | 0.77 | - | - | - |
| HNSC | 1.0 | 0.98 | - | - | - |
| KICH | 1.0 | 0.87 | 0.96 | - | - |
| KIRC | 1.0 | 0.95 | 0.96 | - | 0.9820 |
| KIRP | 1.0 | 0.93 | 0.92 | - | - |
| LIHC | 1.0 | 0.97 | 0.98 | - | - |
| LUAD | 1.0 | 0.95 | 0.96 | $0.9927 \pm 0.0083$ | 0.848 |
| LUSC | 1.0 | 0.91 | 0.88 |  | 0.977 |
| PAAD | 1.0 | 0.97 | 0.95 | - | - |
| PCPG | 1.0 | 1.0 | 1.0 | - | - |
| PRAD | 1.0 | 1.0 | 1.0 | - | - |
| READ | 1.0 | 0.35 | 0.0 | - | - |
| SARC | 1.0 | 0.97 | 0.96 | - | - |
| STAD | 1.0 | 0.96 | - | $0.9943 \pm 0.0039$ | - |
| THCA | 1.0 | 1.0 | 1.0 | $0.9967 \pm 0.0042$ | - |
| THYM | 1.0 | 0.99 | 0.94 | - | - |
| UCEC | 1.0 | 0.96 | 0.96 | - | 0.9640 |
| NORMAL | 1.0 | - | - | - | - |

### 3.4 Component Wise Strength Analysis

In this section, we provide experimental analysis on the strengths of PanClassif. We have performed all the experimental analysis on TCGA data. Firstly, to show the effectiveness of the Anova based feature selection, we have compared it with *χ*^2^ based feature selection and compared the results with Anova based method. Average MCC and accuracy for each classifier were taken using the best 105, 204, 571, 743, 899 genes selected by both methods. A box plot of the MCC for these two methods for binary and multi-class classification task is shown in Fig. 7 (i). As shown in the figure, Anova based feature selection performs significantly better MCC scores for both tasks. ANOVA achieves average MCC of 0.84 and 0.85 compared to those of 0.80 and 0.83 achieved by *χ*^2^ method. (See supplementary table S1, S3, S5, S7 for details.)

**Figure 7:**
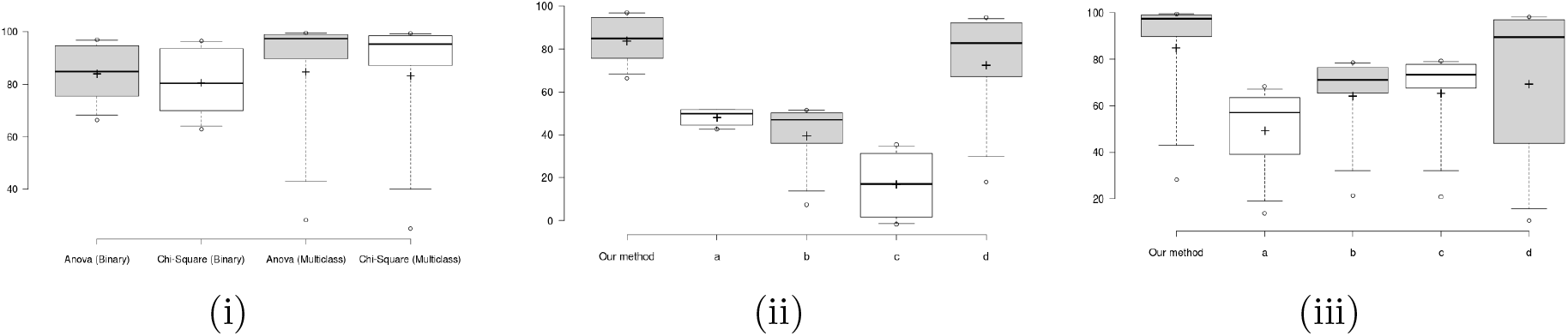
Box plot of MCC scores for (i) different feature selection techniques in binary and multi-class classification tasks, different components of PanClassif on (ii) binary classification task and (iii) multi-class classification task.

To show the effectiveness of smoothing and oversampling steps of PanClassif, we have created 4 variants: (a) under-sampled data with smoothing, (b) over-sampled data without smoothing and, (c) imbalanced data without smoothing (d) imbalanced data with smoothing. Fig. 7 (ii) and (iii) shown box plot of MCC scores for all these variants along with PanClassif run using different models on binary and multi-class classification tasks on TCGA data (please see supplementary table S9, S10, S11 for details).

In the case of undersampling, our train data with 6213 cancer samples were down-sampled by picking 8.20% samples randomly from each cancer types resulting into 509 cancer samples. This is variant ‘a’ in Fig. 7 (ii-iii). This variant achieves average MCC score of 0.48 and 0.49 in binary and multi-class tasks which is very poor compared to those of 0.84 and 0.85 respectively. The other variant ‘b’ in Fig. 7 (ii-iii) was created by oversampling and removing smoothing step. This variant shows the superiority of oversampling over undersampling in multi-class classification by giving MCC values of 0.39 and 0.64 in binary and multi-class respectively.

The variant ‘c’ in Fig. 7 (ii-iii) was generated without over-sampling and without smoothing which shows an average MCC of 0.16 in binary and 0.65 in multi-class classification. However, adding smoothing in imbalance data, dictates its excellence by an average MCC of 0.77 and 0.69 in binary and multi-class classification respectively. This case is represented by the variant ‘d’ in Fig. 7 (ii-iii).

Instead of standardization, we have also performed normalization on TCGA data and found that it gives a performance boost on binary classification. In pan cancer classification we can see that, SVM(kernel=RBF)’s performance increases as we have performed normalization and slightly performance decreasing on Adaboost. Also we can notice that, while doing pan-cancer classification, as we increase the unique no. of genes in our experiment the performance of every classifiers increases when we use normalization on our data (Please see supplementary table S14 and S15 for details). We have included both standardization and normalization mode in our package (check package documentation for details).

### 3.5 Gene Enrichment Analysis

We have further performed gene enrichment analysis with our top 105 genes using the ‘Enrichr web tool’ [33, 34] to validate if the top genes found by PanClassif are associated with cancer cell line. A horizontal bar plot of our top genes against the different cell lines are shown in Fig. 8. The figure shows 6 cancer enriched genes among top 10. They are Pulmonary Sclerosing Hemangioma, Clear-cell metastatic renal cell carcinoma, Conventional (Clear Cell) Renal Cell carcinoma, Invasive Ductal breast Carcinoma, Pancreatic Neoplasm and Kidney Neoplasm’. With this finding, we can conclude that PanClassif is able to identify cancer related best significant genes and that is why all classifiers are performing well.

**Figure 8:**
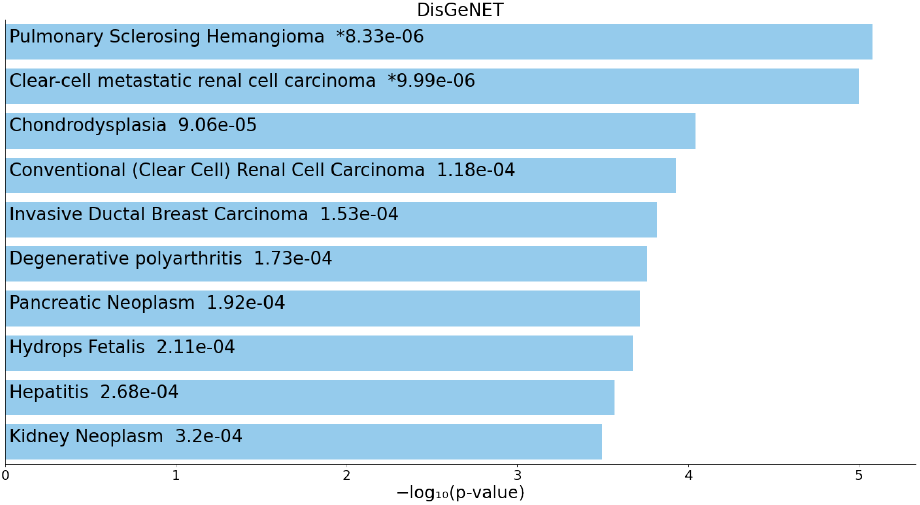
Bar Chart of gene enrichment analysis.

### 3.6 Gene Significance Test

To further test the effectiveness of PanClassif, we have performed gene significance test in two separate experiments. In the first experiment, we have created two datasets: raw data from the TCGA repository and the other one after applying smooting and over-sampling as defined in PanClassif. This time we have selected only 22 genes by selecting only the top gene from each cancer type. After that we have used a neural network classifier to perform multi-class classification on these two datasets. In Table 5, we have shown best genes selected for each cancer and respective sensitivity or recall for each of the dataset. As we can see in Table 5, except CHOL and READ cancer type, PanClassif improves the sensitivity of classifying a cancer type using the top 22 genes. Also note that two cancer types, BRCA and PCPG have already attained a 100% sensitivity using only 22 genes.

**Table 5:** Comparison table of true positive rates between raw data and PanClassif process data using top 22 genes.

| cancer type | top gene | raw data | Pan-Classif | cancer type | top gene | raw data | Pan-Classif |
| --- | --- | --- | --- | --- | --- | --- | --- |
| BLCA | CLEC3A | 37.25 | 89.71 | LUAD | STX10 | 40.97 | 87.5 |
| BRCA | ADAMTS4 | 66.78 | 100 | LUSC | GDR115 | 46.61 | 81.88 |
| CESC | SEH1L | 32.47 | 92.21 | PAAD | CT45A1 | 50.00 | 78.26 |
| CHOL | PROX2 | 16.66 | 0.00 | PCPG | FRMPD4 | 76.92 | 100 |
| COAD | SCARA3 | 47.69 | 86.25 | PRAD | TMIGD2 | 58.74 | 87.68 |
| ESCA | RGMA | 16.98 | 18.36 | READ | LRRC39 | 11.11 | 0.00 |
| HNSC | CA9 | 48.95 | 88.02 | SARC | PTGR2 | 65.33 | 98.48 |
| KICH | DZIP1L | 42.11 | 56.52 | STAD | GCNT7 | 46.55 | 86.61 |
| KIRC | SFPQ | 66.45 | 84.86 | THCA | TLA2G7 | 73.05 | 73.94 |
| KIRP | TEKT5 | 64.36 | 86.42 | THYM | S100B | 79.31 | 56.67 |
| LIHC | ANGPTL5 | 66.39 | 83.01 | UCEC | TRIM16 | 39.65 | 85.71 |

In the second experiment, we have used t-SNE plot on the whole TCGA and GEO datasets. This time we have used 105 top genes which produced satisfactory results in classification. The t-SNE plots for TCGA and GEO multi-class classification task is shown in Fig. 9 for raw data and data transformed using PanClassif. From the fig. 9(b) and 9(d), we can clearly see that the samples is clustered according to their classes in the transformed version using PanClassif. Here it may seems like the samples are decreased in fig. 9(b) but it just put same clustered samples in a compact region. On the other hand, in Fig. 9(a) and 9(c) samples of each classes overlaps one another (no transformation). This will lead to errors in classification.

**Figure 9:**
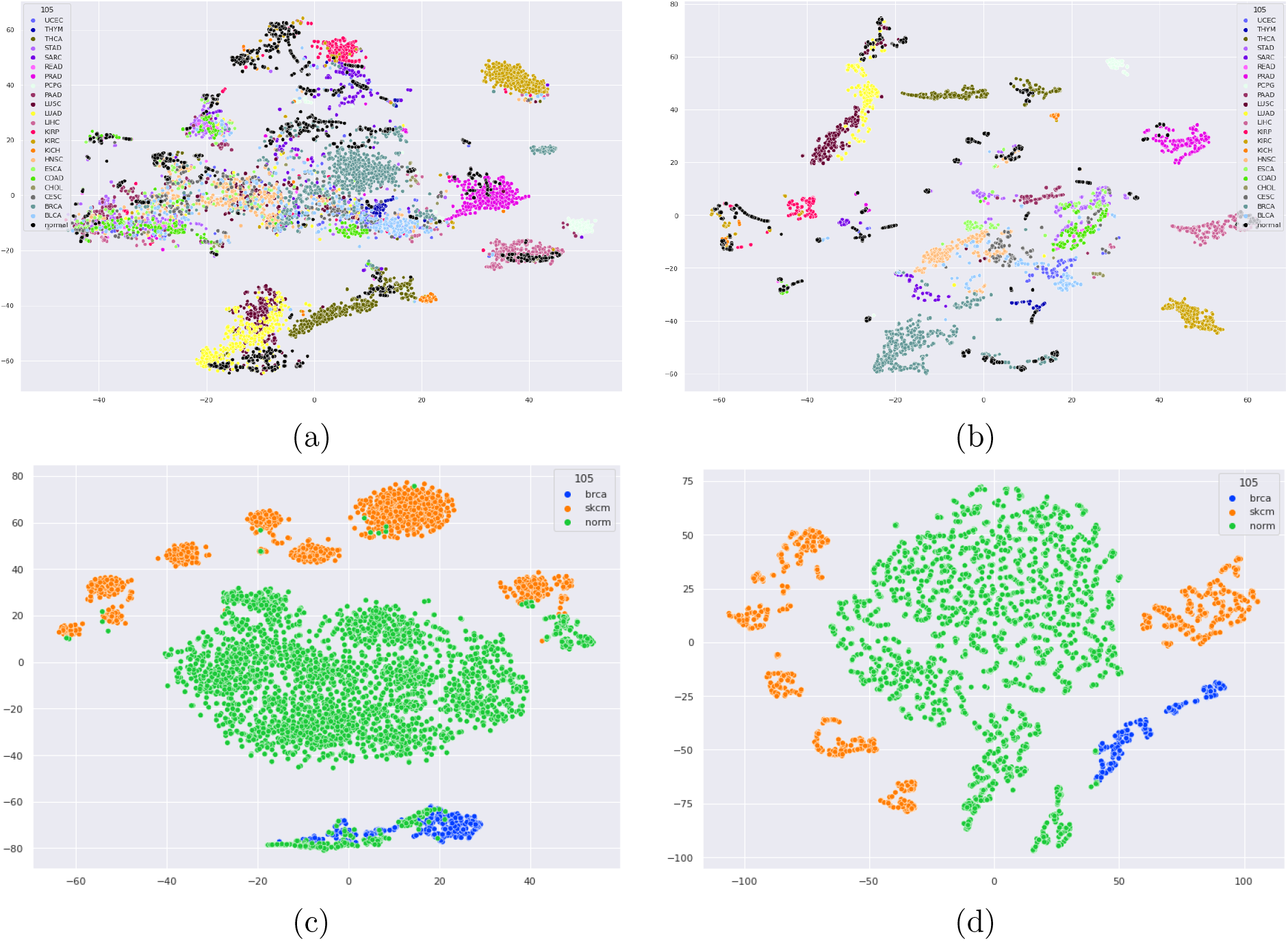
t-SNE of (a) raw TCGA data with no transformation, (b) TCGA’s processed data using PanClassif, (c) raw GEO data with no transformation and (d) GEO processed data using PanClassif

### 3.7 Availability of PanClassif

PanClassif is available as Python package in Python package index (PyPI). The latest version can be installed using `pip install panclassif`. Documentation of PanClassif is also available at https://pypi.org/project/panclassif/#description. PanClassif requires RNA-Seq data as input after smoothing. PanClassif requires a number of folders to be specified as data input and very easy to use. PanClassif codes are available at https://github.com/Zwei-inc/panclassif and the data used in this experiment are uploaded to a open source data repository to ensure reproducibility. The top 105 genes are listed in the supplementary Table S12.

### 3.8 Discussion

PanClassif presents a pipeline to treat TCGA pan cancer RNA-seq samples in such a way that different cancer phenotype can be classified and distinguished very efficiently with a less number of features. After applying PanClassif on TCGA datasets using 105 top genes we have found better distinguishable clusters in comparison to untreated data as shown in Fig. 9. PanClassif achieves consistent performance in most of the machine learning models in terms of accuracy and MCC in TCGA and GEO datasets. For Binary Classification of cancer and normal samples data from TCGA, PanClassif is able to classify more than 99% accurately using both *k*-nearest neighbour and Random Forest with the MCC 0.97 on both models. These results are significantly better than those reported by [9] using 300 genes. However, Adaboost models(decision tree as base classifier) have performed very poorly on TCGA data as the datasets are high dimensional data which might have lead to curse of dimensionality. Due to this reason the calculation complexity of decision tree classifier have increased exponentially with the increasing depth of tree thus ADB perform poorly.

We have further experimented PanClassif with single cell RNA-seq data from Gene Expression Omnibus (GEO) and got the consistent performance. We get MCC 0.98 on *k*-NN, MCC 0.99 on RF and MCC 0.98 on ANN with the GEO data as well, but L-SVM perform poor with GEO datasets and the reason behind that can be skewed data. L-SVM is not a good choice for skewed datasets.

We have performed gene enrichment analysis with top 105 genes and came to know that ‘PTHLH’ gene can disable or hamper cell apoptosis which prevent cancer or tumor cells to die. We have found that if we select best one gene from each cancer type of TCGA using PanClassif still they can correctly classify more than 74% cancer and normal samples. This might be due to the fact that these particular genes are greatly responsible for those cancer phenotype. Further research may reveal there other significance or insight in cancer study. Our method can help cancer study related to pan cancer types.

## 4 Conclusion

In this paper, we propose PanClassif, an effective single cell RNA-seq data based cancer classification method. Experimental results shows the effectiveness of the proposed method on binary cancer prediction and type identification on TCGA and GEO datasets and thus achieving significantly improved results over state-of-the-art method. The gene enrichment analysis and significance tests shows evidence for the overall effectiveness of PanClassif. Finally, a python package is made available for use. We believe, it is possible to extend the PanClassif methodology to general cell classification tasks using single-cell RNA-Seq data. It is important to note that most of the single cell experiments might not show the real expression counts of the genes and it is possible to further improve the results on a wider range of datasets using sophisticated imputing techniques.

## Notes

https://pypi.org/project/panclassif/

### Competing Interest Statement

The authors have declared no competing interest.

## References

[1] M. Margulies, M. Egholm, W. E. Altman, S. Attiya, J. S. Bader, L. A. Bemben, J. Berka, M. S. Braverman, Y. J. Chen, Z. Chen, S. B. Dewell, L. Du, J. M. Fierro, X. V. Gomes, B. C. Godwin, W. He, S. Helgesen, C. H. Ho, C. H. Ho, G. P. Irzyk, S. C. Jando, M. L. Alenquer, T. P. Jarvie, K. B. Jirage, J. B. Kim, J. R. Knight, J. R. Lanza, J. H. Leamon, S. M. Lefkowitz, M. Lei, J. Li, K. L. Lohman, H. Lu, V. B. Makhijani, K. E. McDade, M. P. McKenna, E. W. Myers, E. Nickerson, J. R. Nobile, R. Plant, B. P. Puc, M. T. Ronan, G. T. Roth, G. J. Sarkis, J. F. Simons, J. W. Simpson, M. Srinivasan, K. R. Tartaro, A. Tomasz, K. A. Vogt, G. A. Volkmer, S. H. Wang, Y. Wang, M. P. Weiner, P. Yu, R. F. Begley, and J. M. Rothberg. Genome sequencing in microfabricated high-density picolitre reactors. Nature, 437(7057):376–380, Sep 2005.

[2] Mingye Hong, Shuang Tao, Ling Zhang, Li-Ting Diao, Xuanmei Huang, Shaohui Huang, Shu-Juan Xie, Zhen-Dong Xiao, and Hua Zhang. Rna sequencing: new technologies and applications in cancer research. Journal of Hematology & Oncology, 13(1):166, Dec 2020.

[3] Assieh Saadatpour, Shujing Lai, Guoji Guo, and Guo-Cheng Yuan. Single-cell analysis in cancer genomics. Trends in genetics : TIG, 31(10):576–586, Oct 2015. S0168-9525(15)00140-7[PII].

[4] Gökcen Eraslan, Lukas M. Simon, Maria Mircea, Nikola S. Mueller, and Fabian J. Theis. Single-cell rna-seq denoising using a deep count autoencoder. Nature Communications, 10(1):390, Jan 2019.

[5] Konstantina Kourou, Themis P. Exarchos, Konstantinos P. Exarchos, Michalis V. Karamouzis, and Dimitrios I. Fotiadis. Machine learning applications in cancer prognosis and prediction. Computational and Structural Biotechnology Journal, 13:8–17, 2015.

[6] Y. Li, K. Kang, J. M. Krahn, N. Croutwater, K. Lee, D. M. Umbach, and L. Li. A comprehensive genomic pan-cancer classification using The Cancer Genome Atlas gene expression data. BMC Genomics, 18(1):508, 07 2017.

[7] Y. H. Hsu and D. Si. Cancer Type Prediction and Classification Based on RNA-sequencing Data. Annu Int Conf IEEE Eng Med Biol Soc, 2018:5374–5377, Jul 2018.

[8] Boyu Lyu and Anamul Haque. Deep Learning Based Tumor Type Classification Using Gene Expression Data. In Proceedings of the 2018 ACM International Conference on Bioinformatics, Computational Biology, and Health Informatics, pages 89–96, Washington DC USA, August 2018. ACM.

[9] Bong-Hyun Kim, Kijin Yu, and Peter C W Lee. Cancer classification of single-cell gene expression data by neural network. Bioinformatics, 36(5):1360–1366, 10 2019.

[10] Tianyu Wang, Boyang Li, Craig E. Nelson, and Sheida Nabavi. Comparative analysis of differential gene expression analysis tools for single-cell rna sequencing data. BMC Bioinformatics, 20(1):40, Jan 2019.

[11] Wenbo Guo, Dongfang Wang, Shicheng Wang, Yiran Shan, Changyi Liu, and Jin Gu. scCancer: a package for automated processing of single-cell RNA-seq data in cancer. Briefings in Bioinformatics, 07 2020. bbaa127.

[12] Jasleen K. Grewal, Basile Tessier-Cloutier, Martin Jones, Sitanshu Gakkhar, Yussanne Ma, Richard Moore, Andrew J. Mungall, Yongjun Zhao, Michael D. Taylor, Karen Gelmon, Howard Lim, Daniel Renouf, Janessa Laskin, Marco Marra, Stephen Yip, and Steven J. M. Jones. Application of a Neural Network Whole Transcriptome–Based Pan-Cancer Method for Diagnosis of Primary and Metastatic Cancers. JAMA Network Open, 2(4):e192597, April 2019.

[13] Jose Alquicira-Hernandez, Anuja Sathe, Hanlee P. Ji, Quan Nguyen, and Joseph E. Powell. scpred: accurate supervised method for cell-type classification from single-cell rna-seq data. Genome Biology, 20(1):264, Dec 2019.

[14] Daniele Mercatelli, Forest Ray, and Federico M. Giorgi. Pan-cancer and single-cell modeling of genomic alterations through gene expression. Frontiers in Genetics, 10:671, 2019.

[15] Mafalda Falcão Ferreira, Rui Camacho, and Luís F. Teixeira. Using autoencoders as a weight initialization method on deep neural networks for disease detection. BMC Medical Informatics and Decision Making, 20(S5):141, August 2020.

[16] Nour Eldeen M. Khalifa, Mohamed Hamed N. Taha, Dalia Ezzat Ali, Adam Slowik, and Aboul Ella Hassanien. Artificial Intelligence Technique for Gene Expression by Tumor RNA-Seq Data: A Novel Optimized Deep Learning Approach. IEEE Access, 8:22874–22883, 2020.

[17] Wenpin Hou, Zhicheng Ji, Hongkai Ji, and Stephanie C. Hicks. A systematic evaluation of single-cell rna-sequencing imputation methods. Genome Biology, 21(1):218, Aug 2020.

[18] David van Dijk, Roshan Sharma, Juozas Nainys, Kristina Yim, Pooja Kathail, Ambrose J. Carr, Cassandra Burdziak, Kevin R. Moon, Christine L. Chaffer, Diwakar Pattabiraman, Brian Bierie, Linas Mazutis, Guy Wolf, Smita Krishnaswamy, and Dana Pe’er. Recovering Gene Interactions from Single-Cell Data Using Data Diffusion. Cell, 174(3):716–729.e27, July 2018.

[19] Florian Wagner, Yun Yan, and Itai Yanai. K-nearest neighbor smoothing for highthroughput single-cell rna-seq data. bioRxiv, 2018.

[20] Mo Huang, Jingshu Wang, Eduardo Torre, Hannah Dueck, Sydney Shaffer, Roberto Bonasio, John I. Murray, Arjun Raj, Mingyao Li, and Nancy R. Zhang. SAVER: gene expression recovery for single-cell RNA sequencing. Nature Methods, 15(7):539–542, July 2018.

[21] Jacob C. Kimmel and David R. Kelley. scnym: Semi-supervised adversarial neural networks for single cell classification. bioRxiv, 2020.

[22] Songwei Ge, Haohan Wang, Amir Alavi, Eric Xing, and Ziv Bar-joseph. Supervised adversarial alignment of single-cell rna-seq data. Journal of Computational Biology, 0(0):null, 2020. PMID: 33470876.

[23] Tian Tian, Jie Zhang, Xiang Lin, Zhi Wei, and Hakon Hakonarson. Model-based deep embedding for constrained clustering analysis of single cell rna-seq data. Nature Communications, 12(1):1873, Mar 2021.

[24] Woosung Chung, Hye Hyeon Eum, Hae-Ock Lee, Kyung-Min Lee, Han-Byoel Lee, Kyu-Tae Kim, Han Suk Ryu, Sangmin Kim, Jeong Eon Lee, Yeon Hee Park, Zhengyan Kan, Wonshik Han, and Woong-Yang Park. Single-cell RNA-seq enables comprehensive tumour and immune cell profiling in primary breast cancer. Nature Communications, 8(1):15081, August 2017.

[25] I. Tirosh, B. Izar, S. M. Prakadan, M. H. Wadsworth, D. Treacy, J. J. Trombetta, A. Rotem, C. Rodman, C. Lian, G. Murphy, M. Fallahi-Sichani, K. Dutton-Regester, J.-R. Lin, O. Cohen, P. Shah, D. Lu, A. S. Genshaft, T. K. Hughes, C. G. K. Ziegler, S. W. Kazer, A. Gaillard, K. E. Kolb, A.-C. Villani, C. M. Johannessen, A. Y. Andreev, E. M. Van Allen, M. Bertagnolli, P. K. Sorger, R. J. Sullivan, K. T. Flaherty, D. T. Frederick, J. Jane-Valbuena, C. H. Yoon, O. Rozenblatt-Rosen, A. K. Shalek, A. Regev, and L. A. Garraway. Dissecting the multicellular ecosystem of metastatic melanoma by single-cell RNA-seq. Science, 352(6282):189–196, April 2016.

[26] Bo Li and Colin N Dewey. Rsem: accurate transcript quantification from rna-seq data with or without a reference genome. BMC bioinformatics, 12(1):1–16, 2011.

[27] Ying-Wooi Wan, Genevera I Allen, and Zhandong Liu. Tcga2stat: simple tcga data access for integrated statistical analysis in r. Bioinformatics, 32(6):952–954, 2016.

[28] Nitesh V. Chawla, Kevin W. Bowyer, Lawrence O. Hall, and W. Philip Kegelmeyer. Smote: Synthetic minority over-sampling technique. J. Artif. Int. Res., 16(1):321–357, June 2002.

[29] Corinna Cortes and Vladimir Vapnik. Support-vector networks. Machine Learning, 20(3):273–297, Sep 1995.

[30] Tin Kam Ho. The random subspace method for constructing decision forests. IEEE Transactions on Pattern Analysis and Machine Intelligence, 20(8):832–844, 1998.

[31] Yoav Freund and Robert E Schapire. A decision-theoretic generalization of on-line learning and an application to boosting. Journal of Computer and System Sciences, 55(1):119–139, 1997.

[32] J J Hopfield. Neural networks and physical systems with emergent collective computational abilities. Proceedings of the National Academy of Sciences, 79(8):2554–2558, 1982.

[33] M. V. Kuleshov, M. R. Jones, A. D. Rouillard, N. F. Fernandez, Q. Duan, Z. Wang, S. Koplev, S. L. Jenkins, K. M. Jagodnik, A. Lachmann, M. G. McDermott, C. D. Monteiro, G. W. Gundersen, and A. Ma’ayan. Enrichr: a comprehensive gene set enrichment analysis web server 2016 update. Nucleic Acids Res, 44(W1):W90–97, 07 2016.

[34] E. Y. Chen, C. M. Tan, Y. Kou, Q. Duan, Z. Wang, G. V. Meirelles, N. R. Clark, and A. Ma’ayan. Enrichr: interactive and collaborative HTML5 gene list enrichment analysis tool. BMC Bioinformatics, 14:128, Apr 2013.

