## supplementary_data for "PanClassif: Improving pan cancer classification of single cell RNA-seq gene expression data using machine learning"

**Panclassif**

| No. of | kNN | | RF | | L-SVM | | R-SVM | | ADB | | ANN | |
|---|---|---|---|---|---|---|---|---|---|---|---|---|
| unique genes | MCC | ACC | MCC | ACC | MCC | ACC | MCC | ACC | MCC | ACC | MCC | ACC |
| 105 | 0.97 | 1 | 0.97 | 1 | 0.67 | 0.93 | 0.63 | 0.93 | 0.76 | 0.96 | 0.84 | 0.97 |
| 204 | 0.97 | 1 | 0.96 | 1 | 0.72 | 0.95 | 0.66 | 0.93 | 0.82 | 0.97 | 0.94 | 0.99 |
| 571 | 0.97 | 1 | 0.96 | 1 | 0.74 | 0.95 | 0.67 | 0.93 | 0.81 | 0.97 | 0.92 | 0.99 |
| 743 | 0.96 | 0.99 | 0.97 | 0.99 | 0.78 | 0.96 | 0.67 | 0.93 | 0.81 | 0.97 | 0.83 | 0.98 |
| 899 | 0.98 | 1 | 0.96 | 0.99 | 0.79 | 0.96 | 0.69 | 0.93 | 0.82 | 0.97 | 0.94 | 0.99 |

Table S1: MCC and ACC of binaryclass classification for 6 classifiers using our method on TCGA data.

| No. of | kNN | | RF | | L-SVM | | R-SVM | | ADB | | ANN | |
|---|---|---|---|---|---|---|---|---|---|---|---|---|
| unique genes | MCC | ACC | MCC | ACC | MCC | ACC | MCC | ACC | MCC | ACC | MCC | ACC |
| 105 | 0.98 | 0.99 | 0.998 | 1 | -0.0015 | 0.68 | 0.96 | 0.98 | 1 | 1 | 0.97 | 0.99 |
| 204 | 0.97 | 0.98 | 1 | 1 | 0.66 | 0.84 | 0.96 | 0.98 | 1 | 1 | 0.98 | 0.99 |
| 571 | 0.94 | 0.98 | 1 | 1 | 0.69 | 0.86 | 0.99 | 0.99 | 1 | 1 | 0.98 | 0.99 |
| 743 | 0.92 | 0.96 | 1 | 1 | 0.76 | 0.89 | 0.94 | 0.95 | 1 | 1 | 0.98 | 0.99 |
| 899 | 0.91 | 0.96 | 1 | 1 | 0.78 | 0.9 | 0.94 | 0.95 | 1 | 1 | 0.97 | 0.97 |

Table S2: MCC and ACC of binaryclass classification for 6 classifiers using our method on GEO data.

| No. of | kNN | | RF | | L-SVM | | R-SVM | | ADB | | ANN | |
|---|---|---|---|---|---|---|---|---|---|---|---|---|
| unique genes | MCC | ACC | MCC | ACC | MCC | ACC | MCC | ACC | MCC | ACC | MCC | ACC |
| 105 | 0.995 | 1 | 0.996 | 1 | 0.96 | 0.96 | 0.88 | 0.89 | 0.32 | 0.3 | 0.98 | 0.98 |
| 204 | 0.994 | 1 | 0.994 | 1 | 0.96 | 0.95 | 0.86 | 0.87 | 0.34 | 0.3 | 0.97 | 0.98 |
| 571 | 0.994 | 1 | 0.996 | 1 | 0.96 | 0.97 | 0.87 | 0.88 | 0.25 | 0.19 | 0.98 | 0.98 |
| 743 | 0.993 | 0.99 | 0.997 | 1 | 0.97 | 0.97 | 0.87 | 0.88 | 0.25 | 0.19 | 0.99 | 0.99 |
| 899 | 0.992 | 0.99 | 0.994 | 0.99 | 0.98 | 0.98 | 0.89 | 0.9 | 0.25 | 0.2 | 0.99 | 0.99 |

Table S3: MCC and ACC of multiclass classification for 6 classifiers using our method on TCGA data.

| No. of | kNN | | RF | | L-SVM | | R-SVM | | ADB | | ANN | |
|---|---|---|---|---|---|---|---|---|---|---|---|---|
| unique genes | MCC | ACC | MCC | ACC | MCC | ACC | MCC | ACC | MCC | ACC | MCC | ACC |
| 105 | 0.86 | 0.94 | 1 | 1 | 0 | 0.69 | 0.88 | 0.94 | 0.98 | 0.99 | 0.94 | 0.97 |
| 204 | 0.8433 | 0.92 | 1 | 1 | 0 | 0.7 | 0.89 | 0.95 | 0.99 | 1 | 0.94 | 0.97 |
| 571 | 0.82446 | 0.91 | 1 | 1 | 0 | 0.7 | 0.87 | 0.94 | 0.998 | 1 | 0.96 | 0.98 |
| 743 | 0.806 | 0.9 | 1 | 1 | -0.008 | 0.65 | 0.85 | 0.92 | 1 | 1 | 0.956 | 0.98 |
| 899 | 0.81 | 0.9 | 1 | 1 | -0.074 | 0.6 | 0.84 | 0.92 | 0.95 | 0.98 | 0.96 | 0.99 |

Table S4: MCC and ACC of multiclass classification for 6 classifiers using our method on GEO data.

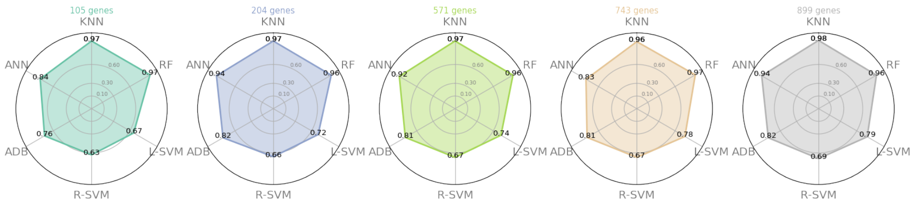

Figure S1: Spider plot for MCC score for the binary classification task in TCGA data using 5 different datasets created selecting different number of genes.

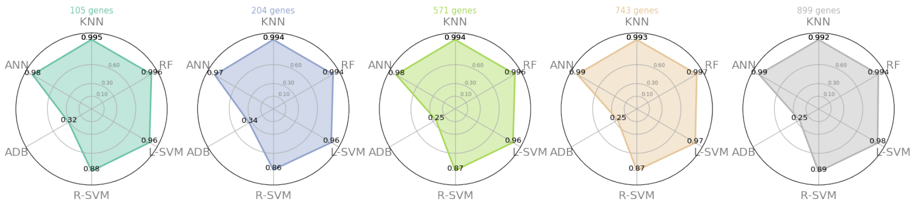

Figure S2: Spider plot for MCC score for the multi-class classification task in TCGA data using 5 different datasets created selecting different number of genes.

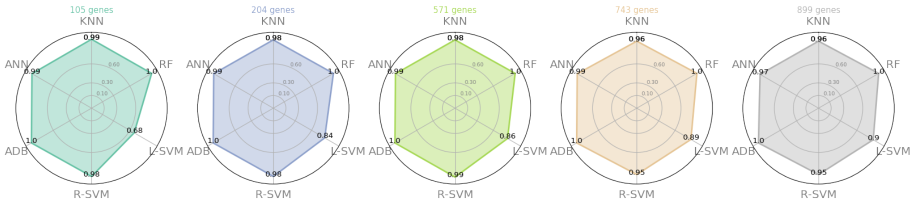

Figure S3: Spider plot for ACC score for the binary classification task in GEO data using 5 different datasets created selecting different number of genes.

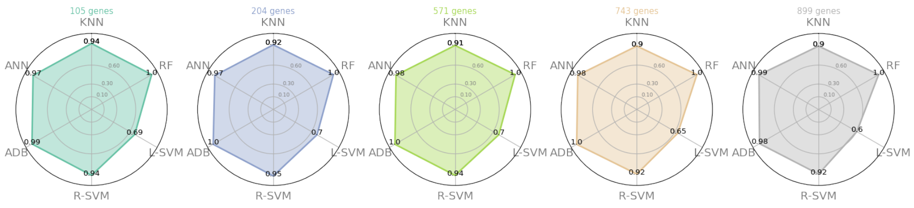

Figure S4: Spider plot for ACC score for the multi-class classification task in GEO data using 5 different datasets created selecting different number of genes.

| No. of unique genes | kNN | | RF | | L-SVM | | R-SVM | | ADB | | ANN | |
|---|---|---|---|---|---|---|---|---|---|---|---|---|
| | MCC | ACC | MCC | ACC | MCC | ACC | MCC | ACC | MCC | ACC | MCC | ACC |
| 69 | 0.98 | 0.98 | 0.991 | 0.99 | 0.9 | 0.9 | 0.74 | 0.75 | 0.17 | 0.2 | 0.96 | 0.97 |
| 109 | 0.99 | 0.99 | 0.991 | 0.99 | 0.92 | 0.93 | 0.84 | 0.85 | 0.09 | 0.17 | 0.97 | 0.97 |
| 136 | 0.99 | 0.99 | 0.995 | 1 | 0.93 | 0.93 | 0.82 | 0.83 | 0.22 | 0.25 | 0.94 | 0.94 |
| 368 | 0.99 | 0.99 | 0.994 | 0.99 | 0.95 | 0.96 | 0.9 | 0.9 | 0.16 | 0.22 | 0.98 | 0.98 |
| 477 | 0.994 | 0.99 | 0.996 | 1 | 0.97 | 0.97 | 0.9 | 0.91 | 0.16 | 0.23 | 0.98 | 0.99 |
| 567 | 0.994 | 0.99 | 0.994 | 0.99 | 0.96 | 0.97 | 0.9 | 0.91 | 0.7 | 0.16 | 0.98 | 0.99 |

Table S7: MCC and ACC of multiclass classification for 6 classifiers using Chi-square as feature selection on TCGA data.

| No. of unique genes | kNN | | RF | | L-SVM | | R-SVM | | ADB | | ANN | |
|---|---|---|---|---|---|---|---|---|---|---|---|---|
| | MCC | ACC | MCC | ACC | MCC | ACC | MCC | ACC | MCC | ACC | MCC | ACC |
| 69 | 0.91 | 0.96 | 1 | 1 | 0 | 0.69 | 0.93 | 0.97 | 0.93 | 0.97 | 0.97 | 0.99 |
| 109 | 0.89 | 0.95 | 1 | 1 | 0 | 0.68 | 0.9 | 0.96 | 0.996 | 1 | 0.95 | 0.98 |
| 136 | 0.86 | 0.93 | 1 | 1 | 0 | 0.68 | 0.88 | 0.95 | 0.99 | 1 | 0.95 | 0.98 |
| 368 | 0.82 | 0.92 | 1 | 1 | 0 | 0.69 | 0.88 | 0.95 | 0.96 | 0.98 | 0.94 | 0.97 |
| 477 | 0.81 | 0.91 | 1 | 1 | 0 | 0.68 | 0.89 | 0.95 | 0.996 | 1 | 0.94 | 0.97 |
| 567 | 0.73 | 0.87 | 1 | 1 | 0 | 0.7 | 0.85 | 0.93 | 0.93 | 0.97 | 0.95 | 0.98 |

Table S8: MCC and ACC of multiclass classification for 6 classifiers using Chi-square as feature selection on GEO data.

| No. of unique genes | kNN | | RF | | L-SVM | | R-SVM | | ADB | | ANN | |
|---|---|---|---|---|---|---|---|---|---|---|---|---|
| | MCC | ACC | MCC | ACC | MCC | ACC | MCC | ACC | MCC | ACC | MCC | ACC |
| 69 | 0.96 | 0.99 | 0.96 | 0.99 | 0.54 | 0.91 | 0.49 | 0.91 | 0.68 | 0.94 | 0.81 | 0.97 |
| 109 | 0.96 | 0.99 | 0.95 | 0.99 | 0.64 | 0.92 | 0.57 | 0.92 | 0.75 | 0.95 | 0.83 | 0.97 |
| 136 | 0.97 | 1 | 0.96 | 0.99 | 0.65 | 0.92 | 0.6 | 0.92 | 0.74 | 0.96 | 0.84 | 0.97 |
| 368 | 0.96 | 0.99 | 0.97 | 0.99 | 0.71 | 0.94 | 0.7 | 0.93 | 0.78 | 0.96 | 0.83 | 0.97 |
| 477 | 0.97 | 1 | 0.98 | 1 | 0.77 | 0.96 | 0.7 | 0.94 | 0.8 | 0.97 | 0.9 | 0.99 |
| 567 | 0.97 | 1 | 0.96 | 0.99 | 0.77 | 0.96 | 0.71 | 0.94 | 0.79 | 0.96 | 0.9 | 0.99 |

Table S5: MCC and ACC of binaryclass classification for 6 classifier using Chi-square as feature selection on TCGA data.

| No. of unique genes | kNN | | RF | | L-SVM | | R-SVM | | ADB | | ANN | |
|---|---|---|---|---|---|---|---|---|---|---|---|---|
| | MCC | ACC | MCC | ACC | MCC | ACC | MCC | ACC | MCC | ACC | MCC | ACC |
| 69 | 0.99 | 1 | 1 | 1 | 0.66 | 0.85 | 0.97 | 0.98 | 1 | 1 | 0.99 | 1 |
| 109 | 0.98 | 0.99 | 1 | 1 | -0.005 | 0.68 | 0.97 | 0.99 | 1 | 1 | 0.98 | 0.99 |
| 136 | 0.97 | 0.99 | 1 | 1 | 0.28 | 0.73 | 0.97 | 0.99 | 1 | 1 | 0.98 | 0.99 |
| 368 | 0.97 | 0.99 | 1 | 1 | 0.73 | 0.89 | 0.97 | 0.99 | 1 | 1 | 0.98 | 0.99 |
| 477 | 0.96 | 0.98 | 1 | 1 | 0.75 | 0.89 | 0.98 | 0.99 | 1 | 1 | 0.99 | 0.99 |
| 567 | 0.95 | 0.98 | 1 | 1 | 0.79 | 0.91 | 0.97 | 0.99 | 1 | 1 | 0.98 | 0.99 |

Table S6: MCC and ACC of binaryclass classification for 6 classifiers using Chi-square as feature selection on GEO data.

| Models | Anova (Binary) | Chi-Square (Binary) | Anova (Multiclass) | Chi-Square (Multiclass) |
|---|---|---|---|---|
| k-NN | 97 | 96.5 | 99.36 | 99.0167 |
| RF | 96.4 | 96.3333 | 99.54 | 99.35 |
| L-SVM | 74 | 68 | 96.6 | 93.8333 |
| R-SVM | 66.4 | 62.8333 | 87.4 | 85 |
| ADB | 80.4 | 75.6667 | 28.2 | 25 |
| NN | 89.4 | 85.1667 | 98.2 | 96.8333 |

Table S9: Feature selection comparison ANOVA vs Chi-Square. Here we put 5 dataset average MCC scores for all the 6 models datasets contains ( 105, 204, 571, 743, 899) genes respectively

| Models | Our method | Under-sampled data with smoothing | Over-sampled data without smoothing | Imbalanceddata without smoothing | Imbalanced data with smoothing |
|---|---|---|---|---|---|
| k-NN | 99.36 | 51.2 | 64.6 | 66.42 | 98.2 |
| RF | 99.54 | 63 | 74.2 | 75.4 | 97.68 |
| L-SVM | 96.6 | 63.8 | 78.6 | 79.2 | 95 |
| R-SVM | 87.4 | 35 | 68.2 | 71.4 | 84 |
| ADB | 28.2 | 13.8 | 21.4 | 20.8 | 30.6 |
| NN | 98.2 | 68.4 | 77.2 | 78.6 | 10.6 |

Table S10: Our method vs 4 variant comparative analysis for multiclass classification. Here we put 5 dataset average MCC scores for all the 6 models. datasets contains ( 105, 204, 571, 743, 899) genes respectively

| Models | Our method | Under-sampled data with smoothing | Over-sampled data without smoothing | Imbalanced data without smoothing | Imbalanced data with smoothing |
|---|---|---|---|---|---|
| k-NN | 97 | 43 | 50.6 | 32.8 | 92.4 |
| RF | 96.4 | 51.8 | 7.4 | -1.68 | 94.8 |
| L-SVM | 74 | 48.6 | 51.6 | 7.2 | 65 |
| R-SVM | 66.4 | 42.6 | 49.4 | -0.12 | 18 |
| ADB | 80.4 | 51.2 | 44.6 | 26.8 | 91.4 |
| NN | 89.4 | 51.8 | 33.2 | 35.4 | 74 |

Table S11: Our method vs 4 variant comparative analysis for Binary Classification. Here we put 5 dataset average MCC scores for all the 6 models. datasets contains ( 105, 204, 571, 743, 899) genes respectively

| | | | | |
|---|---|---|---|---|
| SCARA3 | FLRT2 | RIMS3 | SEH1L | PTGR2 |
| TAAR6 | TEKT5 | RDM1 | CLIC4 | ANO4 |
| ZBTB12 | RIBC2 | PLAC8 | LRRC39 | FIG4 |
| PEX13 | GGT5 | DZIP1L | GLP1R | CFDP1 |
| WDR77 | RASSF8 | C1QTNF6 | OXT | PTHLH |
| SCN8A | CKAP5 | KL | FHIT | DPRX |
| ITGBL1 | C1ORF189 | SFTPB | ECHS1 | GREM1 |
| RELL2 | TGIF2LX | CMTM4 | ADAMTS12 | ANGPTL4 |
| COL9A3 | LCN2 | KCNIP4 | FXR2 | NEK10 |
| FCN1 | TPP2 | TRIM60 | EMB | PI15 |
| TFAP2A | MAP3K14 | ASNS | OR5AN1 | ANGPTL5 |
| C2ORF86 | APOBEC3B | EMP1 | CA9 | SPRY1 |
| FRMPD4 | UBIAD1 | KRT12 | CD300LG | GCNT1 |
| FGL2 | PDE1C | MMP9 | GPR115 | ARHGDIG |
| STX10 | LPAR2 | TGFBR1 | CD34 | BARX1 |
| ABAT | GATA5 | TMEM175 | VENTX | FAM48B1 |
| ATP2C1 | GPR180 | RGMA | LONRF1 | CST4 |
| C1ORF113 | LOC284233 | TMIGD2 | ANGEL2 | ADAMTS4 |
| CLEC3A | PLA2G7 | KAT2A | KCTD3 | GPR132 |
| GNG5 | CT45A1 | SFPQ | PROX2 | RNASE8 |
| PEBP4 | NPPC | TMEM219 | S100B | GCNT7 |

Table S12: Top 105 unique gene name list using our method.

| Cancer Name | Top gene | Raw data | | | PanClassif | | |
|---|---|---|---|---|---|---|---|
| | | Test samples | Accurately predicted | TPR | Test samples | Accurately predicted | TPR |
| BLCA | CLEC3A | 102 | 38 | 37.25 | 107 | 96 | 89.72 |
| BRCA | ADAMTS4 | 292 | 195 | 66.78 | 303 | 303 | 100 |
| CESC | SEH1L | 77 | 25 | 32.46 | 77 | 71 | 92.21 |
| CHOL | PROX2 | 12 | 2 | 16.66 | 11 | 0 | 0 |
| COAD | SCARA3 | 65 | 31 | 47.69 | 80 | 69 | 86.25 |
| ESCA | RGMA | 53 | 9 | 16.98 | 49 | 9 | 18.36734694 |
| HNSC | CA9 | 143 | 70 | 48.95 | 142 | 125 | 88.02 |
| KICH | DZIP1L | 19 | 8 | 42.10 | 23 | 13 | 56.52 |
| KIRC | SFPQ | 158 | 105 | 66.45 | 152 | 129 | 84.86 |
| KIRP | TEKT5 | 87 | 56 | 64.36 | 81 | 70 | 86.42 |
| LIHC | ANGPTL5 | 122 | 81 | 66.39 | 106 | 88 | 83.01 |
| LUAD | STX10 | 144 | 59 | 40.97 | 144 | 126 | 87.5 |
| LUSC | GDR115 | 133 | 62 | 46.61 | 138 | 113 | 81.88 |
| PAAD | CT45A1 | 44 | 22 | 50 | 46 | 36 | 78.26 |
| PCPG | FRMPD4 | 39 | 30 | 76.92 | 47 | 47 | 100 |
| PRAD | TMIGD2 | 143 | 84 | 58.74 | 138 | 121 | 87.68 |
| READ | LRRC39 | 27 | 3 | 11.11 | 26 | 0 | 0 |
| SARC | PTGR2 | 75 | 49 | 65.33 | 66 | 65 | 98.48 |
| STAD | GCNT7 | 116 | 54 | 46.55 | 112 | 97 | 86.60 |
| THCA | TLA2G7 | 141 | 103 | 73.05 | 142 | 105 | 73.94 |
| THYM | S100B | 29 | 23 | 79.31 | 30 | 17 | 56.66 |
| UCEC | TRIM16 | 58 | 23 | 39.65 | 49 | 42 | 85.71 |

Table S13: Top 22 gene's significance by accuracy for every individual cancer for raw and PanClassif processed data using multiclass classification.

| No. of unique genes | **kNN** | | RF | | L-SVM | | R-SVM | | ADB | | ANN | |
|---|---|---|---|---|---|---|---|---|---|---|---|---|
| | MCC | ACC | MCC | ACC | MCC | ACC | MCC | ACC | MCC | ACC | MCC | ACC |
| 105 | 0.9905 | 1 | 1 | 1 | 0.91 | 0.99 | 0.98 | 1 | 0.97 | 0.99 | 0.997 | 1 |
| 204 | 0.996 | 1 | 1 | 1 | 0.96 | 0.99 | 1 | 1 | 1 | 1 | 0.996 | 1 |
| 571 | 1 | 1 | 1 | 1 | 0.96 | 0.99 | 1 | 1 | 1 | 1 | 1 | 1 |
| 743 | 0.99 | 0.99 | 1 | 1 | 0.97 | 1 | 0.97 | 0.99 | 0.98 | 1 | 0.97 | 1 |
| 899 | 1 | 1 | 1 | 1 | 0.99 | 1 | 0.99 | 0.99 | 1 | 1 | 0.99 | 1 |

Table S14: MCC and ACC of binaryclass classification for 6 classifiers using normalization on TCGA data.

| No. of unique genes | **kNN** | | RF | | L-SVM | | R-SVM | | ADB | | ANN | |
|---|---|---|---|---|---|---|---|---|---|---|---|---|
| | MCC | ACC | MCC | ACC | MCC | ACC | MCC | ACC | MCC | ACC | MCC | ACC |
| 105 | 0.95 | 0.99 | 0.95 | 0.98 | 0.79 | 0.8 | 0.85 | 0.91 | 0.07 | 0.15 | 0.94 | 0.97 |
| 204 | 0.96 | 0.99 | 0.98 | 0.99 | 0.87 | 0.88 | 0.89 | 0.93 | 0.25 | 0.29 | 0.96 | 0.98 |
| 571 | 0.98 | 1 | 0.997 | 1 | 0.96 | 0.98 | 0.88 | 0.94 | 0.21 | 0.23 | 0.95 | 0.99 |
| 743 | 0.97 | 0.99 | 0.99 | 1 | 0.98 | 0.99 | 0.94 | 0.95 | 0.19 | 0.2 | 0.98 | 1 |
| 899 | 0.99 | 1 | 1 | 1 | 0.99 | 1 | 0.94 | 0.95 | 0.23 | 0.21 | 0.99 | 1 |

Table S15: MCC and ACC of multiclass classification for 6 classifiers using normalization on TCGA data.

| | Binary Classification | | Multiclass Classification | |
|---|---|---|---|---|
| Models | With k-NN smoothing | With SAVER | With k-NN smoothing | With SAVER |
| Random Forest | 0.97 | 0.67 | 0.995 | 0.75 |
| KNN | 0.97 | 0.73 | 0.996 | 0.68 |
| L-SVM | 0.72 | 0.53 | 0.96 | 0.77 |
| R-SVM | 0.63 | 0.66 | 0.88 | 0.72 |
| AdaBoost | 0.76 | 0.69 | 0.32 | 0.31 |

Table S16: Comparison of imputation methods based on MCC over TCGA dataset.
